# Alarmone ((p)ppGpp) signalling tunes *Arabidopsis thaliana* nuclear gene expression to shape the balance of plant immune outcomes

**DOI:** 10.64898/2026.07.17.739251

**Authors:** Federico E Aballay, Ivan J León Sanchez, Florencia S Rodriguez, Camila Benelli, Pedro J Salaberry, Denise Scuffi, Carlos García-Mata, Rocío S Tognacca, Nicolas M Cecchini, Ben Field, Ezequiel Petrillo

**Author notes:** Corresponding authors: Federico E. Aballay < >, Nicolas M. Cecchini < >, Ben Field < > & Ezequiel Petrillo < >.

## Abstract

RSH enzymes (RelA/SpoT homologs) synthesize ppGpp (guanosine tetra-/pentaphosphate) in plastids, regulating organelle function. More importantly, whether ppGpp acts as a simple rheostat on immune output or determines which mode of defence a plant deploys remains poorly understood. Using *Arabidopsis thaliana* lines with high (RSH3OX) or null (*rshq*) ppGpp levels, we show that ppGpp controls nuclear-encoded defence genes in salicylic acid (SA) and jasmonic acid (JA) metabolism, including hormone-inactivating enzymes in RSH3OX and MeSA-to-SA conversion genes in *rshq*. Conversely, pathogen infection induces the ppGpp synthases RSH2 and RSH3, with induction preceding defence gene activation and requiring SA biosynthesis and a functional Type III secretion system. RSH3OX plants are hypersusceptible to *Pseudomonas syringae* pv. *tomato* DC3000, show impaired pattern-triggered immunity, and exhibit an accelerated decline in photosynthetic efficiency, while *rshq* plants show enhanced resistance to *P. syringae* but increased susceptibility to the necrotrophic fungus *Botrytis cinerea*. Using an inducible line to acutely elevate ppGpp, we confirm this susceptibility is directly caused by ppGpp levels, not a chronic developmental consequence of RSH3OX. Together, these results indicate that ppGpp governs the mode of immune execution rather than simply scaling its magnitude, acting as a chloroplast-based checkpoint linking organellar status to the nuclear defence transcriptome.

## Introduction

Plants have evolved sophisticated mechanisms to integrate environmental information and coordinate proper developmental and stress responses. Oxygenic photosynthesis, which fixes carbon from atmospheric CO₂ and releases O₂ as a byproduct, is the ultimate foundation of life on Earth. Food chains, and our ability to breathe, ultimately depend on photosynthetic organisms that harness sunlight to produce their own energy. Oxygenic photosynthesis, which uses water as the primary electron donor, is conducted by cyanobacteria and their endosymbiotic descendants, the chloroplasts of plant cells (Miyagishima, 2023). Over evolutionary time, the endosymbiont has relinquished most of its genome: plastids retain only ∼5% of the ortholog genes found in cyanobacteria, with the nuclear genome encoding the majority of the ∼3000 protein subunits needed for chloroplast function (Martin *et al.,* 2002; Cullis *et al.,* 2009). Optimal photosynthesis therefore depends on the coordinated assembly of protein complexes encoded across different cellular compartments (Adir *et al.,* 2003). This coordination requires strict regulation and high plasticity, ensuring that chloroplast activity adapts to environmental changes. Nuclear–chloroplast communication is mediated by bidirectional signalling: anterograde signals (nucleus to organelle) regulate plastid gene expression, while retrograde signals (organelle to nucleus) adjust nuclear transcription according to plastid status (Pfannschmidt *et al.,* 2003). Today, chloroplasts are recognized not only as sites of photosynthesis but also as environmental sensors that transmit retrograde signals to the nucleus, modulating transcript levels, RNA processing, and translation in response to light and cellular conditions (Jung *et al.,* 2010; Kubaczka *et al*., 2024; Petrillo *et al.,* 2014; Puthiyaveetil *et al.,* 2009). Plastids thereby fine-tune nuclear gene expression in response to developmental cues and other internal and external signals (Chan *et al*., 2016; Nomura *et al*., 2012).

Plant immunity is a multilayered system that relies, mainly, on two branches. i) pattern-triggered immunity (PTI) that is activated due to the sensing of pathogen-associated molecular patterns (PAMPs) such as flagellin, lipopolysaccharides or elongation factor Tu (Ef-Tu). These PAMPs are recognized by pattern recognition receptors (PRRs) at the plasma membrane (Macho and Zipfel, 2014; Jones *et al*., 2024); ii) effector-triggered immunity (ETI), which is activated by pathogen effectors delivered into host cells through diverse mechanisms, such as the bacterial type III secretion system (T3SS) (O’Malley *et al*., 2021), or by effector-induced modifications of host targets. Effectors or the alterations they cause can be recognized directly or indirectly by intracellular nucleotide-binding leucine-rich repeat receptors (NLRs) (Jones *et al*., 2016; Jones *et al*., 2024). PTI and ETI responses vary in the magnitude and duration but result in similar downstream molecular events, such as mitogen-activated protein kinase (MAPK) activation, oxidative burst, ion influx, Ca^2+^ signaling, increased biosynthesis of plant defense hormones, and transcriptional reprogramming (Ngou *et al*., 2021; Yu *et al*., 2024). Moreover, plant immunity is tightly regulated by hormonal pathways such as salicylic acid (SA) and jasmonic acid (JA). SA and JA regulate both local and systemic immune responses. SA contributes to defense-gene activation and the regulation of hypersensitive cell death and plays a central role in systemic acquired resistance (SAR). In some plant species, its volatile derivative methyl salicylate (MeSA) can function as a mobile signal during SAR, whereas jasmonates have also been implicated in long-distance immune signalling (Park *et al*., 2007; Liu *et al*., 2011; Truman *et al*., 2007).

Plastids act as central hubs for plant environmental adaptation. These organelles couple phytohormone synthesis with retrograde signalling to coordinate nuclear gene expression, serving not merely as sensors but as active executors (Chan *et al.,* 2015). Driven by photosynthesis, their reactive oxygen species (ROS)-mediated communication network bridges stress perception and defence, orchestrating functions from metabolic reprogramming to direct pathogen elimination (Caplan *et al.,* 2015; Noctor *et al.,* 2018; Littlejohn, *et al*., 2020; Rui *et al*., 2025). As a corollary, disruption of chloroplast function often results in altered defence responses (Serrano *et al*., 2016). However, despite all the accumulated knowledge, the molecular nature of plastid-to-nucleus signals remains poorly understood. We hypothesize that signalling mechanisms present in bacteria may have been co-opted during plant evolution to serve for nuclear gene expression control. Among these, the signalling molecule guanosine tetraphosphate/pentaphosphate ((p)ppGpp), hereafter referred to as ppGpp, has emerged as a key regulator of chloroplast function and plant metabolism (Mehrez *et al*., 2023), and could also be relevant to regulate nuclear gene expression (Sugliani *et al*., 2016; Romand *et al*., 2022; Inazu *et al*., 2024). In bacteria, ppGpp, originally characterized as the "alarmone" in the stringent response, accumulates upon exposure to various stresses to inhibit global transcription and certain enzymatic activities to overcome stress (Cashel & Gallant, 1969; Liu *et al*., 2016; Bange *et al.,* 2021). In plants, ppGpp is synthesized in chloroplasts by RelA/SpoT Homolog (RSH) enzymes and modulates chloroplast size, number and gene expression, particularly in response to light transitions and nutrient availability (Sugliani *et al.,* 2016; Ono *et al*., 2020; D’Alessandro *et al*., 2024).

*Arabidopsis thaliana* encodes four plastid-localized RSH enzymes in the nuclear genome: RSH1, RSH2, RSH3 and cRSH. RSH1 has an active hydrolase domain and an inactive synthase domain. In contrast, cRSH has an active synthase domain and an inactive hydrolase domain. In addition, RSH2 and RSH3 are paralogs with functional hydrolase and synthase domains. Plastids are also intimately connected to the salicylic acid (SA) and jasmonic acid (JA) signalling pathways, which makes the plastidial production of ppGpp potentially relevant to immunity and hormone signalling (Pieterse *et al*., 2012; Wildermuth *et al*., 2001; Wasternack & Hause, 2013; Romand *et al.,* 2022). While SA accumulation activates systemic acquired resistance (SAR) and is essential for defence against hemi-biotrophic pathogens like *Pseudomonas syringae,* JA and its bioactive conjugate JA-isoleucine (JA-Ile) mainly mediate resistance to necrotrophic fungi and herbivorous insects (Glazebrook, 2005). These pathways exhibit extensive crosstalk, often functioning antagonistically to fine-tune defence outputs based on the nature of the threat (Thaler *et al*., 2012). The homeostasis of both hormones is tightly regulated through biosynthetic, catabolic, and conjugation pathways. For example, methylation of SA by BENZOIC ACID/SA CARBOXYL METHYLTRANSFERASE 1 (BSMT1) to produce methyl salicylate (MeSA) and glycosylation by UGT74F2 to produce into SA glucose ester (SGE) represent major inactivation routes. In contrast, demethylation of MeSA by METHYL ESTERASE (MES) enzymes regenerates active SA (Vlot *et al*., 2009; Klessig *et al*., 2018). Similarly, JA-Ile is deactivated by oxidation (catalyzed by CYP94 enzymes) and conjugation reactions (Koo *et al*., 2011).

Emerging evidence links ppGpp to plant defence responses. JA treatment induces ppGpp accumulation in *Pisum sativum*, whereas ppGpp overaccumulation reduces SA levels in *Arabidopsis thaliana*, including in the context of viral infection (Takahashi *et al*., 2004; Abdelkefi *et al*., 2018). Beyond this hormonal crosstalk, ppGpp itself accumulates upon infection with the pathogen *Pseudomonas syringae* pv. *tomato* DC3000 (*Pst*), and similarly, upon flg22 treatment, indicating that pathogen-associated molecular pattern (PAMP) receptor-mediated signalling may control ppGpp levels (Qiu *et al*. 2023). Conversely, ppGpp-deficient plants accumulate higher levels of both SA and JA, suggesting a broader role for ppGpp in hormonal homeostasis (Inazu *et al*., 2024). Thus, whether ppGpp accumulation is ultimately beneficial or detrimental to plant immunity, and how it shapes nuclear defence gene expression and pathogen resistance, remains poorly understood.

In this study, we investigated the role of ppGpp in regulating *A. thaliana* immunity using transcriptomic profiling and functional pathogen assays. We show that ppGpp levels control the expression of a broad spectrum of nuclear-encoded defence genes, particularly those involved in SA and JA metabolism. ppGpp overaccumulation (RSH3OX) upregulates genes encoding SA and JA inactivating enzymes, while plants with null levels of ppGpp (*rshq*) show an upregulation of genes involved in SA activation (MeSA demethylation). Consistent with these transcriptional changes, plants with high ppGpp levels exhibit hypersusceptibility to the hemi-biotrophic pathogen *Pseudomonas syringae* pv. *tomato* DC3000 (*Pst*), whereas those without ppGpp display enhanced resistance. Conversely, ppGpp-null plants are hypersusceptible to the necrotrophic fungus *Botrytis cinerea*, consistent with elevated SA antagonizing JA-mediated defence. Temporal analysis of *RSH* gene expression during pathogen infection reveals that *RSH2* and *RSH3*, that encode the major ppGpp synthases during pathogen responses, are rapidly induced, suggesting that ppGpp accumulation is an early event in the defence response. Together, our findings establish ppGpp as a chloroplast-based signal that determines the mode of immune execution a plant deploys, rather than simply scaling its magnitude, with opposing consequences for resistance to pathogens of different lifestyles.

## Results

### ppGpp regulates nuclear-encoded immunity-related genes linked to JA and SA signalling

ppGpp’s known functions in plants are related to growth, photosynthesis regulation and chloroplast control (Mehrez *et al*., 2023). However, the effects of the alarmone on the plant nuclear transcriptome are poorly characterized. To understand how nuclear gene expression is modulated by ppGpp levels in normal growth conditions, we evaluated the transcriptomes of two-week old *A. thaliana* seedlings comparing a wild-type line (Col-0) to an *RSH3* overexpression line (RSH3OX), with high levels of ppGpp, and an *RSH* quadruple mutant (*rshq*), with null levels of ppGpp. These lines were previously characterized by Sugliani *et al*. (2016) and Inazu *et al*. (2024), respectively. Seedlings were grown in MS-agar plates in constant light for two weeks. This allowed plastid development while reducing circadian oscillations (Petrillo *et al*., 2014). We then transferred half of the plates to darkness and kept the other half in light for 48 hours to set up contrasting light and dark conditions. Interestingly, the vast majority of the genes affected by light/dark in the WT are equally affected in the lines with high or null levels of ppGpp (Figure 1A). Hence, disrupting alarmone levels in seedlings does not severely affect light or dark-shaped transcriptomes. However, we found a plethora of nuclear genes that are differentially expressed (DE) in the ppGpp-altered lines, in light and in dark conditions, which could be divided into four main clusters (Figure 1B): cluster 1, genes upregulated by elevated ppGpp levels; cluster 2, genes whose expression is downregulated in both RSH3OX and *rshq* plants relative to wild-type (Col- 0); cluster 3, genes upregulated by low/null ppGpp levels; and cluster 4, genes upregulated by both high and null ppGpp levels. GO term analysis of biological processes for DEGs in both lines revealed enrichment in genes related to jasmonic acid (JA) responses and oxylipin biosynthesis. Additional enriched terms included ’response to wounding’, ’negative regulation of defence response to insect’, ’triterpenoid metabolic process’, and ’ethylene-activated signalling pathway’ (Figure 1C). Since these results indicate that the expression of defence-related genes is altered in plants with contrasting ppGpp levels, and given the well-established antagonism between JA and salicylic acid (SA) signalling in plant defence, we next focused on the transcriptional profiles of defence markers alongside representative SA and JA marker genes. In RSH3OX plants, where ppGpp is elevated, we observe potent induction of stress-responsive genes, most notably *BSMT1* and *N-ACETYLTRANSFERASE ACTIVITY 1 (NATA1*), indicating amplified metabolic reprogramming toward hormone inactivation pathway rather than canonical defence activation (Figure 1D). Conversely, *rshq* plants exhibit striking and selective upregulation of specific Type 1 Plant Defensins (*PDF1.2*, *1.2C*, and *1.3*). While classical SA-marker genes such as *PR1* and *PR2* showed modest increases in *rshq* line, the strong *BSMT1* response in RSH3OX plants underscores a shift toward inactive MeSA. Moreover, *rshq* plants exhibit a higher expression of *PREPIP3*, which encodes a pro-peptide that, once cleaved, binds to cell surface receptor RLK7 to enhance plant stress tolerance (Zhou *et al.,* 2022). These results were validated in independent experiments by RT-qPCR (Figure 1E). Taken altogether, these results show ppGpp regulates plant defence signalling.

**Figure 1.**
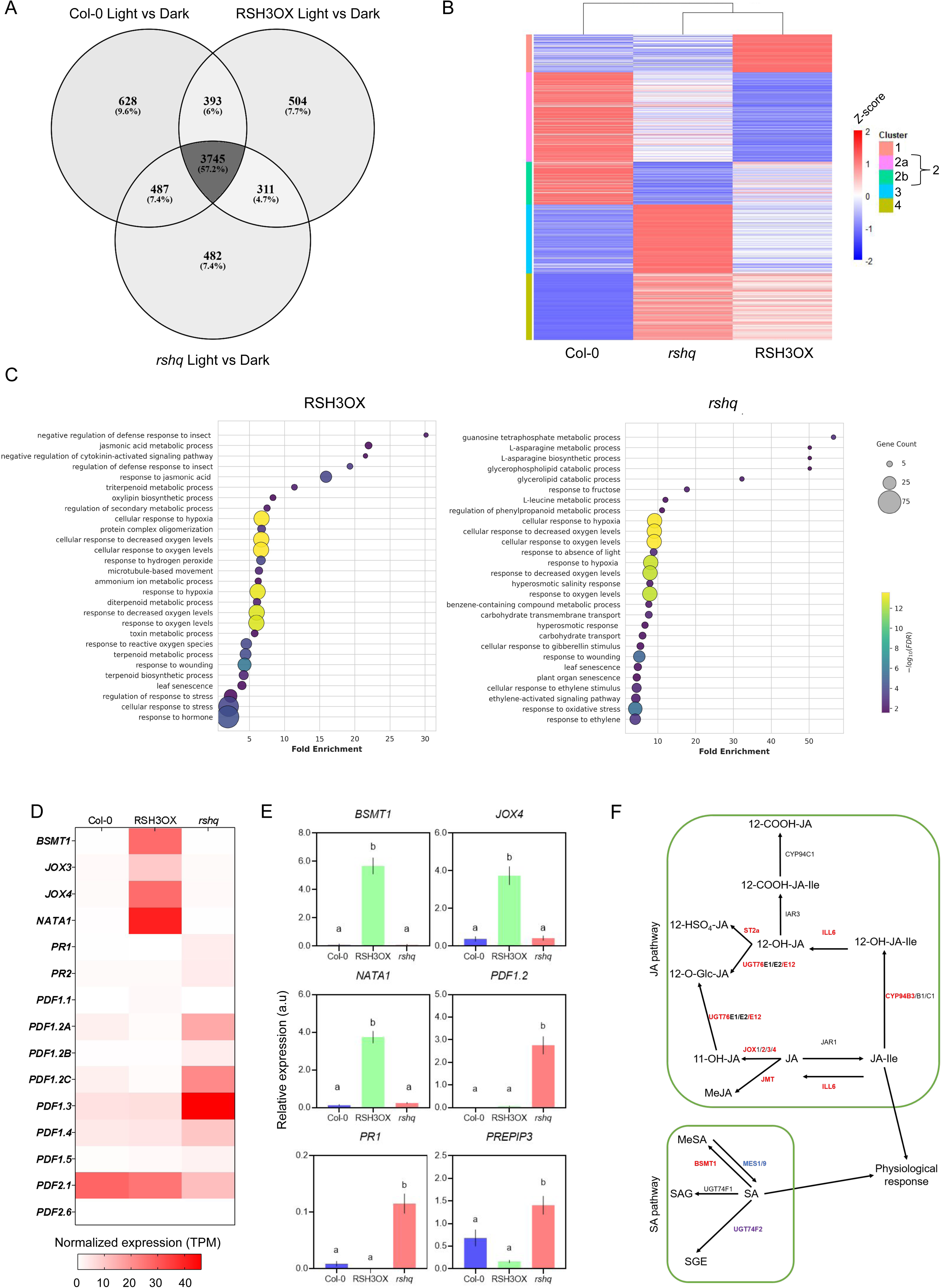
ppGpp-dependent transcriptional reprogramming in *Arabidopsis thaliana*. (A) ppGpp does not substantially affect light and dark-related transcriptomes. Venn diagram showing most of the affected genes in light versus dark in the WT plants (Col-0) are also affected in ppGpp null (*rshq* mutant) or high level (RSH3OX) lines. Venn diagrams were constructed with DEG with |Log2(FC)| > 0.1 and adj. p-value < 0.05. (B) K-means clustering of transcriptional profiling (Z-scores of the means) of whole 14-day-old *Arabidopsis thaliana* seedlings from ppGpp-overaccumulating (RSH3OX) and ppGpp-null (*rshq*) plants in light conditions. Hierarchical clustering of the different genotypes is presented as a dendrogram. (C) Gene Ontology (GO) enrichment analysis of differentially expressed genes (relative to Col-0) shows that RSH3OX plants are enriched in hormone- and stress-related processes, including jasmonic acid metabolism, while *rshq* mutants are enriched in ethylene and oxidative stress responses. Data are shown in bubble plot. Bubble size indicates gene count and color represents statistical significance (-log (FDR)). Data have been previously filtered by FDR ≤ 0.05. (D) Heatmap of selected defense-related genes illustrates impact of ppGpp on salicylic acid (SA) and jasmonic acid (JA) pathways. SA markers (*PR1, PR2*) remain low in RSH3OX, whereas JA oxidases (*JOX3, JOX4*) and other metabolic genes (*BSMT1, NATA1*) are induced. In contrast, *rshq* mutants show enhanced expression of JA-responsive defensins (*PDF* genes, notably *PDF1.2* and *PDF1.3*). This gene set was chosen to exemplify ppGpp’s dual control over SA and JA signaling, highlighting canonical markers, pathway regulators. (E) RT-qPCR validation of representative genes from light data (relative to housekeeping gene *PP2A*). Statistical differences are indicated by distinct letters above bars. Together, these data demonstrate that ppGpp acts as a central integrator of hormone crosstalk, fine-tuning SA and JA outputs to balance metabolic reprogramming with immune defense. Different letters indicate significant differences among genotypes (p < 0.05, one-way ANOVA with Fischer post-hoc test). (F) JA and SA signaling related genes are regulated by ppGpp levels. While high ppGpp levels tend to affect JA signalling, low ppGpp specifically acts in the SA signalling pathway. Red: upregulated in RSH3OX; blue: upregulated in *rshq*; Violet: downregulated in *rshq*. Only genes with |Log2(FC)| ≥ 1 are considered shown. MeJA: Methyl Jasmonate; JA-Ile: Jasmonoyl isoleucine; MeSA: Methyl salicylate; SAG: Salicylic acid glucoside; SGE: Salicylate glucose ester; 11-OH-JA: 11-hydroxyjasmonic acid; 12-COOH-JA: 12-carboxyjasmonic acid; 12-OH-JA-Ile: 12-hydroxyjasmonoyl isoleucine; 12-COOH-JA-Ile: 12-carboxyjasmonoyl isoleucine; 12-HSO-JA: 12-sulfatejasmonic acid; 12-O-Glc-JA: 12-glucoside jasmonate ester.

Beyond SA, ppGpp also shapes jasmonate metabolism, with overaccumulation in RSH3OX upregulating a suite of JA-Ile-deactivating enzymes: *ILL6, UGT76E1, ST2A, CYP94B3, JMT* and *JOX*s (Table S1 and Figure 1F). Notably, no reciprocal upregulation of JA-Ile biosynthesis or signalling genes was detected in *rshq* plants, suggesting that ppGpp influences the JA pathway primarily through activation of JA conjugation metabolism when ppGpp levels are high. Furthermore, *rshq* plants exhibit higher expression of *MES1* and *MES9*, both encoding methyl salicylate esterases that convert MeSA to SA, while simultaneously showing downregulated expression of *UGT74F2*, which encodes a glucoside-ester transferase responsible for converting SA into its storage form, salicylate glucose ester (SGE). Taken together, these expression changes strongly suggest that high ppGpp levels might favour inactive yet volatile MeSA accumulation at the expense of free SA, whereas low ppGpp levels promote free SA accumulation by enhancing MeSA-to-SA conversion and reducing SA storage capacity. Also, in RSH3OX plants, our transcriptomic data suggest a possible shift toward volatile, inactive or storage forms of JA, such as hydroxylated JA.

### Pathogen perception and infection induce *RSH2* and *RSH3* expression

Since the expression of defence-related genes is altered in plants with contrasting ppGpp levels, we next asked whether pathogen and elicitor challenge reciprocally regulate the expression of *RSH* genes, establishing a feedback-loop that could control ppGpp levels (see Figure 2A for the roles of each RSH protein in alarmone accumulation). Pathogen-induced upregulation of *RSH2* and *RSH3* has previously been reported in flood-inoculated *Arabidopsis* seedlings challenged with *Pst* or flg22 (Qiu *et al*., 2023). We therefore asked whether this transcriptional response extends to adult plants inoculated by leaf infiltration and whether it generalizes to additional elicitors and pathogens. Analysis of publicly available transcriptomic data revealed that *RSH2* and *RSH3* transcript levels are upregulated in leaves upon both *Pst* infection and treatment with chitin (Makechemu *et al*., 2025), a fungal cell wall-derived elicitor, while *RSH1* and *cRSH* remained largely unresponsive (Supplementary Figure S1A). A modest but consistent upregulation of *RSH2* was also detected in roots in response to chitin, suggesting that elicitor-triggered induction of ppGpp synthesis genes is not restricted to aerial tissues. These results are further confirmed by analysing *Pst*-infected adult leaves, where *RSH2* and *RSH3* are upregulated in *Pst-*treated samples compared to the mock control, whilst *RSH1* and *cRSH* are not affected (Supplementary Figure S1B). Additionally, by re-analyzing publicly available transcriptomic data of *A. thaliana* leaves infected with *Botrytis cinerea* (Wei *et al.,* 2024), we found that *RSH2* and *RSH3* expression is upregulated over a 96 h time course following spray inoculation of whole plants (Supplementary Figure S1C). Single-cell transcriptomic re-analysis (Zhu *et al*., 2023) confirmed that, upon *Pst* infection, *RSH3* is broadly upregulated across all cell types in the leaf (Supplementary Figure S2A), while within the mesophyll, spanning cell states from immune-related to susceptibility-linked, as defined by Zhu and colleagues (Supplementary Figure S2B-C), both *RSH2* and *RSH3* are induced (Supplementary Figure S2D). This suggests that the induction of ppGpp synthesis capacity, at least from the gene expression regulation perspective, is a specific and reproducible feature of the leaf transcriptional response to microbial infection.

**Figure 2.**
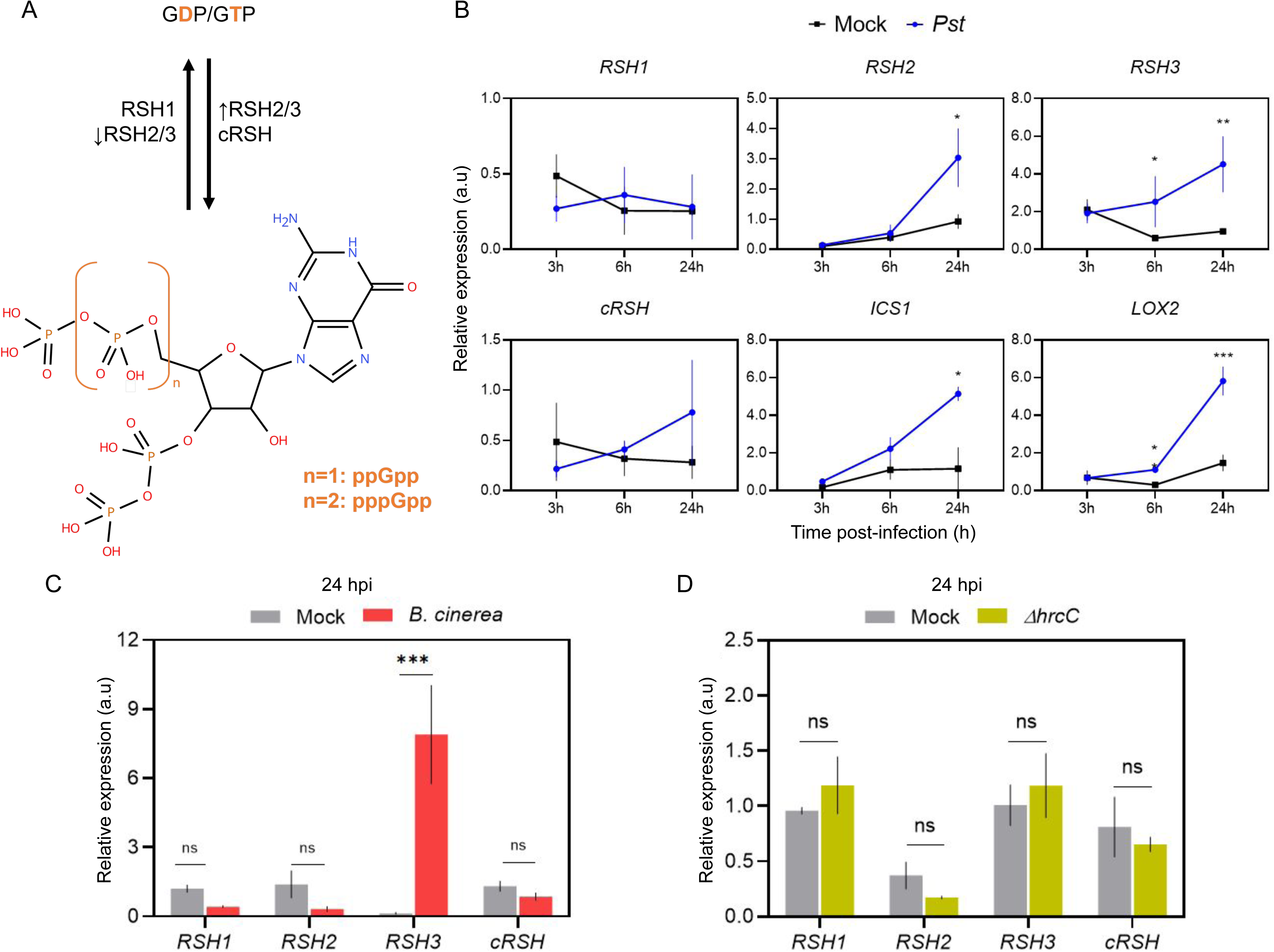
Virulent pathogen infection induces ppGpp biosynthesis genes. **(A)** Schematic representation of (p)ppGpp metabolism in plants. RSH1, RSH2 and RSH3 hydrolyze ppGpp into GDP or GTP and pyrophosphate (PPi, not shown), while RSH2, RSH3, and cRSH can synthesize ppGpp from GDP or GTP and ATP(not shown), releasing AMP. The chemical structure of (p)ppGpp is shown below. If GDP is used as substrate, ppGpp is produced. If GTP is used as substrate, ppGpp is produced. For the case of RSH2 and RSH3, up arrow indicates synthase activity and down arrow indicates hydrolase activity. **(B)** RT-qPCR analysis of *RSH1*, *RSH2*, *RSH3*, *cRSH*, *ISOCHORISMATE SYNTHASE 1* (*ICS1*), and *LIPOXYGENASE 2* (*LOX2*) transcript levels in *A*. *thaliana* leaves at 3, 6, and 24 hours after inoculation with *Pst* (blue, OD_600_= 0.0005) or mock solution (black). Expression values are shown in arbitrary units (a.u.) relative to reference gene *EF-1A*. Data are presented as mean ± SEM of four biological replicates. Asterisks indicate statistically significant differences between treatments within each genotype (p < 0.05, two way ANOVA with post-hoc test). **(C)** RT-qPCR analysis of *RSH1, RSH2, RSH3* and *cRSH* transcript levels in *A. thaliana leaves at* 24 h post inoculation with liquid PD (mock, gray) or *Botrytis cinerea* (10^6^ spores/mL, yellow). Expression values are shown in arbitrary units (a.u.) relative to reference gene *UBQ10*. Data are presented as mean ± SEM of three biological replicates. Asterisks indicate statistically significant differences between mock and *Botrytis cinerea* treatments within each genotype (p < 0.05, two way ANOVA with post-hoc test); ns: not significant. **(D)** RT-qPCR analysis of *RSH1, RSH2, RSH3* and *cRSH* transcript levels in *A. thaliana leaves at* 24 h post infiltration with T3SS-deficient strain Δ*hrcC (*OD_600_=0.01) (yellow) or MgCl_2_ 10 mM (mock, grey). Expression values are shown in arbitrary units (a.u.) relative to reference gene *EF-1A*. Data are presented as mean ± SEM of three biological replicates. Asterisks indicate statistically significant differences between mock and Δ*hrcC* within each genotype (p < 0.05, two way ANOVA with post-hoc test); ns: not significant.

To characterize the temporal dynamics of the induction of ppGpp metabolism and hormone biosynthesis genes, we monitored *RSH* transcript levels by RT-qPCR at 3, 6, and 24 hours after inoculation with *Pst*, alongside the SA biosynthesis marker *ISOCHORISMATE SYNTHASE 1* (*ICS1*), the JA biosynthesis marker *LIPOXYGENASE 2* (*LOX2*), and the defence output marker *PATHOGENESIS-RELATED GENE 1* (*PR1*) (Figure 2B and Supplementary Figure S3). While *RSH1* and *cRSH* transcript levels remained stable throughout the time course, *RSH2* and *RSH3* are progressively induced upon *Pst* infection, with levels rising between 6 and 24 hours post-inoculation. In contrast, *PR1*, *ICS1*, and *LOX2* showed little or no induction at 3 or 6 hours, with strong upregulation only becoming evident at 24 hours post-inoculation (Figure 2B). These results indicate that *RSH2* and *RSH3* induction precedes, or occurs concomitantly with, the transcriptional activation of defence-related genes and hormone biosynthesis pathways, consistent with ppGpp accumulation acting upstream of the defence response rather than as a downstream consequence of it.

In addition, when adult leaves are challenged with *Botrytis cinerea,* a necrotrophic pathogen, we observed that there is a strong induction of *RSH3* expression at 24 hpi, with *RSH1*, *RSH2*, and *cRSH* unaffected (Figure 2C). This contrasts with the more gradual, *RSH2*-dominant induction observed in the public *B. cinerea* spray-inoculation time course (Wei *et al*., 2024; Supplementary Figure S1C), likely reflecting the distinct infection dynamics of a droplet challenge on detached leaves in our system versus a lower-dose, whole-plant spray inoculation followed over a longer time course (Wei *et al.,* 2024). Together, these results indicate that *RSH* transcript levels increase in response to both hemi-biotrophic and necrotrophic pathogens, though the specific *RSH* paralog engaged may depend on infection route and dose. Full induction of *RSH2* and *RSH3* expression required a functional T3SS, as the T3SS-deficient *Pst* mutants *ΔhrpA* and *ΔhrcC* failed to trigger this response (Figure 2D and Supplementary Figure S4). This requirement does not necessarily indicate that a specific effector directly activates *RSH2/RSH3* transcription; it may instead reflect reduced bacterial colonization, diminished chloroplast perturbation, or a less sustained hormonal signal in the absence of T3SS-mediated virulence.

### Induction of *RSH2* and *RSH3* depends on the SA biosynthetic pathway

Having established that infection induces *RSH2* and *RSH3*, we next asked what signal drives this induction. Since pathogen infection triggers hormone signalling in the plant host, we decided to determine whether the expression of *RSH* genes is controlled by hormone signalling pathways. We found that exogenous addition of SA or MeJA for 6 hours slightly downregulates the expression of *RSH1* while not affecting the expression of *RSH*2, *RSH3* and *cRSH* (Supplementary Figure S5A). These results indicate that neither SA nor JA signalling is sufficient for the induction of *RSH2* and *RSH3* gene expression. In addition, we infected ICS1-lacking plants (*sid2*) and *jar1* plants, defective in SA and JA signalling respectively, with *Pst* and analyzed *RSH* transcript levels by RT-qPCR. Notably, *RSH2* and *RSH3* induction was lost in *sid2* but not *jar1* plants, indicating that pathogen induction of *RSH* expression depends on SA but not JA (Supplementary Figure S5B). These results suggest that SA is necessary for inducing *RSH2/3* expression upon pathogen infection. Since exogenous SA alone does not induce *RSH2* or *RSH3* (Supplementary Figure S5A), the requirement for ICS1 during pathogen infection may reflect a role for the SA biosynthetic pathway, or a metabolite thereof, rather than canonical SA signalling.

### ppGpp regulates defence responses against *Pseudomonas syringae* pv. *tomato* DC3000

We found that ppGpp regulates the expression of defence-related genes even in the absence of pathogens (Figure 1), and that pathogen infection reciprocally induces the expression of the genes coding for RSH2 and RSH3, which synthesize ppGpp, suggesting that alarmone levels are in turn modulated during infection. To directly characterize the role of the alarmone in defence responses, we performed pathogen infection assays on RSH3OX and *rshq* lines (Figure 3A). At the lowest inoculum concentration of *Pseudomonas syringae* pv. *tomato* DC3000 (OD₆₀₀ = 0.0001), RSH3OX plants supported significantly greater bacterial growth than wild-type plants (Col-0), indicating enhanced susceptibility, whereas *rshq* plants behaved similarly to the wild type. At the highest inoculum concentration (OD₆₀₀ = 0.001), bacterial growth was comparable between RSH3OX and wild-type plants. In contrast, *rshq* plants supported significantly lower bacterial growth under these conditions, suggesting that the absence of ppGpp enhances resistance at higher pathogen inoculum levels.

**Figure 3.**
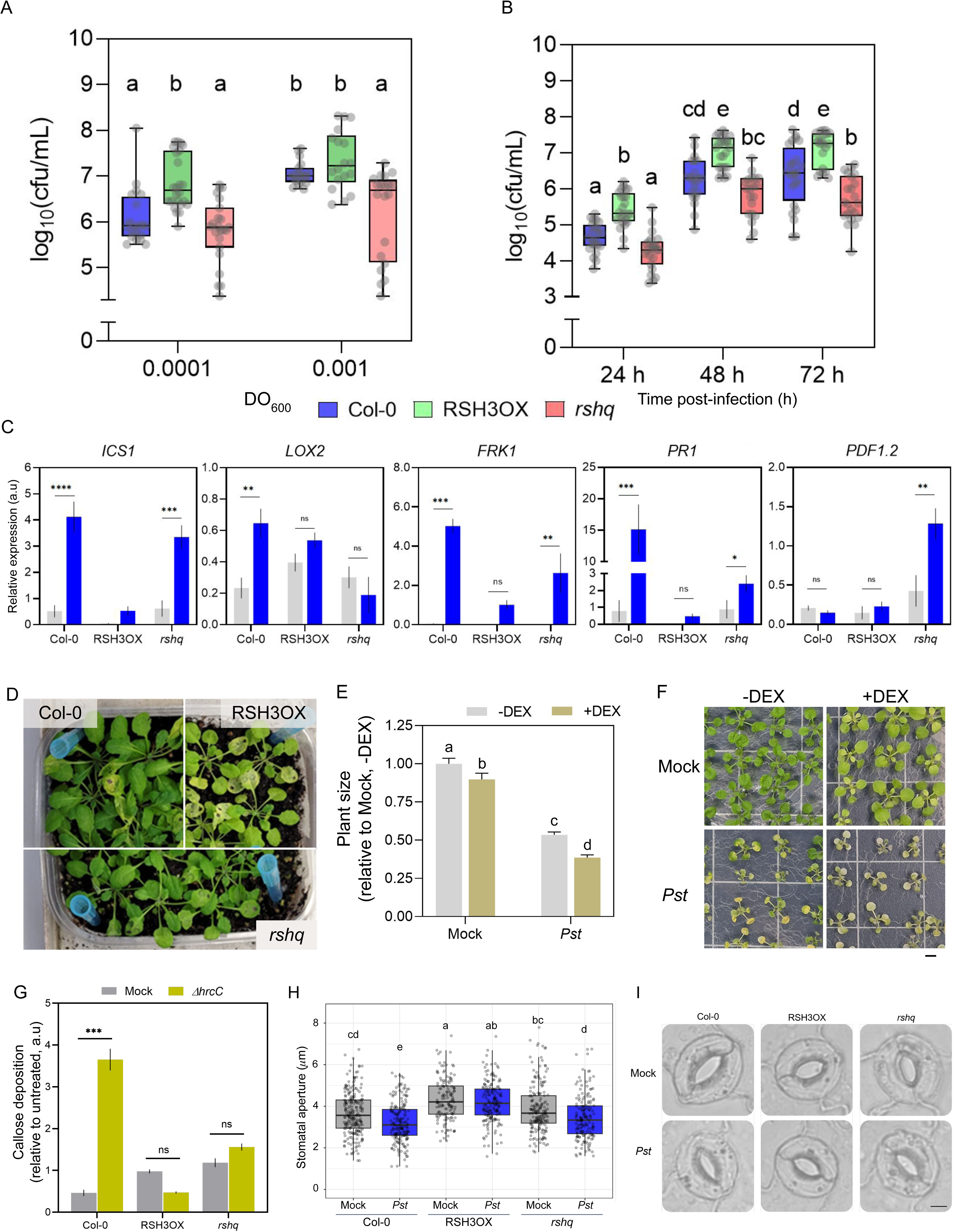
ppGpp levels determine the outcome of *Pseudomonas syringae* pv. *tomato* DC3000 infection. **(A)** Bacterial growth quantification in leaves of Col-0, RSH3OX, and *rshq* plants at 3 days post-inoculation (dpi) with *Pst* DC3000 at two inoculum concentrations (OD₆₀₀ = 0.0001 and 0.001). Bacterial titers are expressed as log₁₀ colony-forming units per mL (cfu/mL). Different letters indicate statistically significant differences between genotypes within each inoculum concentration (p < 0.05, one-way ANOVA with post-hoc test). **(B)** Bacterial growth dynamics in leaves of Col-0, RSH3OX, and *rshq* plants at 24, 48, and 72 hours post-infection (hpi) with *Pst* DC3000 (OD₆₀₀ = 0.0001). Bacterial titers are expressed as log₁₀ (cfu/mL). Different letters indicate statistically significant differences between genotypes at each time point (p < 0.05, one-way ANOVA with post-hoc test). **(C)** RT-qPCR analysis of the SA biosynthesis marker *ISOCHORISMATE SYNTHASE 1* (*ICS1*), the JA biosynthesis gene *LIPOXYGENASE 2* (*LOX2*), the pattern-triggered immunity (PTI) marker *FRK1,* the SA-responsive defense marker *PATHOGENESIS-RELATED 1* (*PR1*), and the JA-responsive marker *PLANT DEFENSIN 1.2A* (*PDF1.2*) in leaves of Col-0, RSH3OX, and *rshq* plants under mock or *Pst* DC3000 infection conditions. Expression values are shown in arbitrary units (a.u.) relative to *EF1A*. Data are presented as mean ± SEM of biological replicates. **(D)** Representative photographs of Col-0, RSH3OX, and *rshq* plants at 3 dpi with virulent *Pst* DC3000 (OD₆₀₀ = 0.0001), illustrating the contrasting infection phenotypes across genotypes. **(E)** Measurement of plant size of infected SYN plants 120 hpi normalized to non-induced (-DEX) and mock-infiltrated plants. Different letters indicate statistically significant differences between treatments (p < 0.05, two-way ANOVA with post-hoc test). **(F)** Representative images of DEX-treated and Pst-infected SYN plants measured in **(E)**. Scale bar: 5 mm. **(G)** Callose deposition in leaves of Col-0, RSH3OX, and *rshq* plants at 24 hpi after mock or Δ*hrcC* inoculation, quantified by aniline blue staining followed by and expressed relative to untreated controls. Asterisks indicate statistically significant differences between mock and Δ*hrcC* within each genotype (p < 0.05, two way ANOVA with post-hoc test); ns: not significant. **(H)** Stomatal aperture quantification on epidermal peels from 5-week old plants. Epidermal peels were preincubated in opening buffer for 3h under light and subsequently exposed to *Pst*. Values are shown in micrometers (*μ*m) and represented in a box plot. Grey boxes represent epidermal peels exposed to mock treatment and blue boxes represent epidermal peels exposed to *Pst*. Individual points represent each measurement. Data are derived from three independent experiments. Different letters indicate significant differences among treatments (Tukey’s method, p-value < 0.05) **(I)** Representative bright field microscopic image of stomata of epidermal peels from both conditions measured in **(F).** Scale bar: 5 μm.

To resolve the temporal dynamics of these phenotypes, we quantified bacterial growth at 24, 48, and 72 hours post-infection (hpi). RSH3OX plants exhibited significantly higher bacterial loads as early as 24 hpi, revealing an accelerated and early bacterial proliferation. In contrast, the shift from wild-type-like susceptibility to enhanced resistance in *rshq* plants was most evident at 72 hpi (Figure 3B), suggesting that ppGpp depletion progressively restricts bacterial proliferation. Altered resistance profiles were only observed in plants infected by wild-type *Pst* and not with the non-pathogenic *ΔhrcC* strain (Supplementary Figure S6), suggesting that T3SS-dependent virulence activity is required to reveal the contrasting phenotypes of the ppGpp-altered lines.

To begin dissecting the molecular basis of these phenotypes, we analysed the expression of defence marker genes in mock- and *Pst*-inoculated plants. At 24hpi with *Pst*, Col-0 plants mount a robust transcriptional induction of the PTI marker *FRK1*, the SA biosynthesis gene *ICS1*, the SA-response marker *PR1*, and the JA biosynthesis gene *LOX2.* The JA signalling indicator *PDF1.2* was not induced. In contrast, RSH3OX plants showed no significant induction of the SA signalling-related genes *ICS1* and *PR1*, nor of the PTI marker *FRK1*, upon *Pst* infection (Figure 3C). *rshq* plants, on the other hand, showed a broadly similar induction pattern to Col-0 (strong induction of *ICS1* and intermediate induction of *PR1* and *FRK1*), with two notable exceptions: *LOX2* induction was reduced to an extent comparable with RSH3OX, and *PDF1.2*, which was not induced in Col-0, was strongly triggered in *rshq* (Figure 3C). Together, these results suggest that elevated ppGpp levels actively suppress canonical immune gene activation, providing a molecular basis for the hypersusceptibility of RSH3OX plants to *Pst* DC3000. The altered profile of defence gene induction in *rshq* plants, combined with their enhanced bacterial growth restriction at 72 hpi, suggests that additional mechanisms may contribute to *Pst* resistance in the absence of ppGpp. Consistent with the bacterial counts and defense-gene expression, the constitutive ppGpp lines also differed in visible disease symptoms: RSH3OX plants showed more severe damage upon *Pst* infection than the wild type, while *rshq* plants were comparatively protected (Figure 3D). However, these phenotypes arise in plants with altered ppGpp throughout their whole life, so the visible damage could be a chronic developmental consequence of the lines rather than a direct effect of ppGpp. To separate these possibilities, we tested whether an acute rise in plastidial ppGpp is sufficient to worsen pathogen-induced damage. We used the SYN transgenic line which expresses a chloroplast-targeted bacterial ppGpp synthase domain from RelA under a dexamethasone (DEX)-inducible promoter, and accumulates high ppGpp levels upon induction (Sugliani *et al*., 2016). DEX-induction of SYN alone caused a modest but significant reduction in plant size relative to uninduced controls under mock conditions (Figure 3E). Upon *Pst* infection, DEX-induced plants showed markedly more severe disease symptoms, including pronounced chlorosis and stunted growth, than infected non-induced controls (Figure 3F), and quantificatification confirmed that plant size was further and significantly reduced in DEX-induced and *Pst-*infected compared to infected plants without DEX induction (Figure 3E). This demonstrates that an acute increase in ppGpp is sufficient to exacerbate pathogen-induced damage even in an otherwise wild-type genetic background, supporting a direct and causal role for ppGpp in shaping the outcome of infection.

### ppGpp homeostasis modulates specific PTI outputs

Given that RSH3OX plants support higher bacterial growth and exhibit downregulated expression of pattern-triggered immunity (PTI) markers like *NCRK* (Supplementary Figure S7A), a gene involved in plasmodesmal callose deposition (Vu *et al*., 2023), we asked whether this susceptibility extended to cell wall-based defences. We therefore assessed callose production in all three genotypes following inoculation with the T3SS-deficient *Pst* mutant Δ*hrcC*, which triggers PTI without delivering effectors into plant cells (Yuan *et al*., 1996; Hauck *et al*., 2003; Peppino Margutti *et al*., 2025). In Col-0, Δ*hrcC* inoculation triggered robust and significant callose deposition relative to mock-treated plants, whereas neither RSH3OX nor *rshq* plants mounted a significant callose response (Figure 3G), indicating that both lines are impaired in this branch of PTI regardless of their opposing susceptibility phenotypes. Nevertheless, growth of the *ΔhrcC* strain was similar across genotypes (Supplementary Figure S6). Thus, the reduced callose response was not sufficient to alter bacterial growth under these conditions, likely because additional PTI mechanisms redundantly restrict this non-virulent strain. These findings also indicate that the enhanced growth of virulent *Pst* DC3000 in RSH3OX plants cannot be explained solely by defective callose deposition.

Since both genotypes show defective callose deposition despite their opposing bacterial growth phenotypes, callose cannot account for this difference. We therefore asked whether stomatal immunity, a second major PTI output that restricts bacterial entry (Pantaleno *et al.,* 2021), might explain the divergence in bacterial growth. These PTI responses are themselves regulated by T3SS effectors that either promote stomatal opening to favour bacterial entry, or closure to favour bacterial growth through water soaking (Kim, et al., 2005; O’Malley, et al., 2021; Roussin- Léveillée, et al., 2022). Given the impaired PTI responses already observed in RSH3OX plants and that *RSH3* is expressed in guard cells upon bacterial infection (Figure 2A-B and Supplementary Figure S2A), we examined whether ppGpp levels influence pathogen-triggered stomatal behaviour. At the transcriptional level, we found that both RSH3OX and *rshq* exhibit differential regulation of genes involved in stomatal aperture and closure (*BTB-A2.3, IQM1, PME53, RZFP34, ABCG40, ALMT5 and FLP*; Supplementary Figure S6B) (Cai *et al.,* 2005; Kang, *et al*., 2010; Zhou *et al*., 2012; Ding *et al* 2015; Wu *et al*., 2022; Doireau *et al*., 2023). Hence, we tested the role of ppGpp levels in stomatal closure by measuring aperture in epidermal peels exposed to *Pst* for 1 hour. In Col-0 and *rshq*, stomatal aperture was reduced in response to *Pst*, whereas in RSH3OX it was not affected. Moreover, RSH3OX plants showed wider stomatal apertures than the wild type, even in mock conditions, suggesting that high ppGpp levels can affect stomatal processes beyond the immune response to *Pst* (Figure 3H and 3I).

### T3SS effectors are required for the regulation of ppGpp-dependent photosynthesis efficiency

Photosynthetic parameters such as maximum photosystem II quantum yield (Fv/Fm) and Non-Photochemical quenching (NPQ) are regulated by both pathogen infection and ppGpp (Kopczewski *et al*., 2020; Pérez-Bueno *et al*., 2019; Sugliani *et al*., 2016; Romand *et al*., 2025). To explore this link, we infected RSH3OX and *rshq* plants with *Pst* WT and monitored Fv/Fm up to 72 hpi. In line with previous reports, upon *Pst* DC3000 infection Col-0 plants exhibited a downregulation in Fv/Fm at 72 hpi (Figure 4A-C). In contrast, Fv/Fm was markedly reduced in RSH3OX plants as early as 48 hpi during *Pst* DC3000 infection. Strikingly, the *rshq* lines did not exhibit any significant changes in photosynthesis efficiency during infection. These results indicate that ppGpp is necessary for the pathogen-induced decline in photosynthetic efficiency. The T3SS is required to downregulate photosynthesis efficiency during pathogen infection in wild type plants (de Torres Zabala *et al.,* 2015). We therefore evaluated changes in Fv/Fm during infection with the *Pst* Δ*hrcC* strain that lacks the T3SS. As expected, Col-0 and *rshq* plants did not exhibit significant changes in Fv/Fm. RSH3OX plants also showed no significant decline in Fv/Fm relative to mock, although chlorophyll fluorescence imaging revealed localized areas of reduced signal in some *ΔhrcC*-infected RSH3OX leaves (Figure 4A-C), suggesting a local effect. Together, these results indicate that the ppGpp-dependent decline in photosynthetic efficiency in the context of infection requires a functional T3SS. Consistent with this interpretation, treatment with flg22 alone did not significantly alter Fv/Fm in any genotype, including RSH3OX (Supplementary Figure S8), confirming that PAMP perception alone, independently of ppGpp status, is not sufficient to trigger the decline in photosynthetic efficiency, at least under the conditions tested.

**Figure 4.**
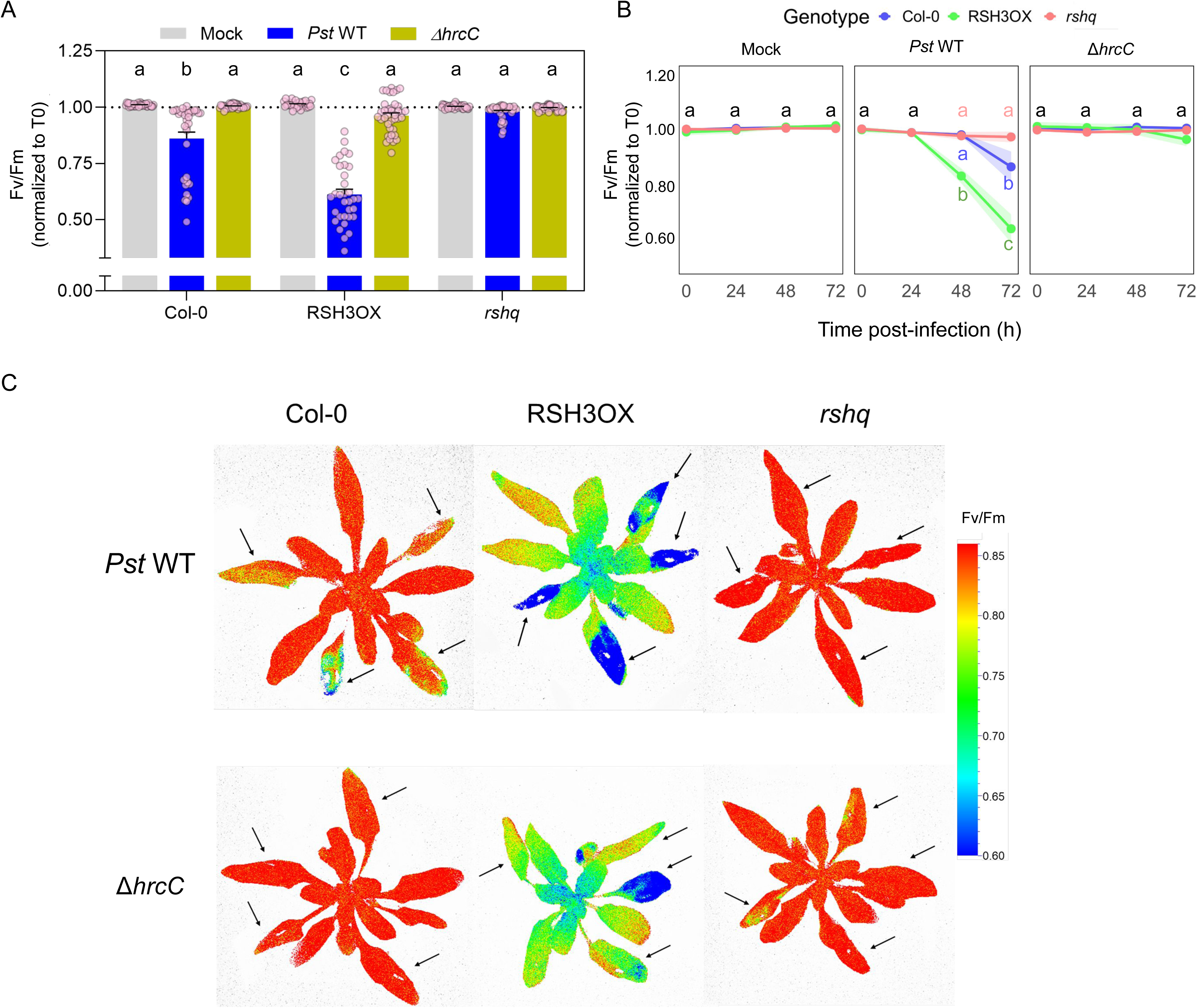
*Pst* DC3000 T3SS-dependent effectors reduce photosynthetic efficiency in a ppGpp-dependent manner. **(A)** Photosynthesis efficiency (Fv/Fm) of adult *A. thaliana* plants infected with mock (MgCl_2_ 10 mM), *Pst* WT or Δ*hrcC* at 72 hpi. Data are shown in arbitrary units (a.u.) relative to 0h post-infection (T0). Data are presented as mean ± SEM. Different letters indicate statistical significant differences among genotypes and treatments (p<0.05, two-way ANOVA with post-hoc test). **(B)** Kinetics of photosynthesis efficiency during *Pst* WT or Δ*hrcC* infection up to 72 hpi. Data are shown in arbitrary units (a.u.) relative to 0h post-infection (T0). Data are presented in line plots as mean ± SEM. Ribbons represent the SEM. Different letters indicate statistical significant differences among time points (p<0.05). Black letters indicate same statistical result among all groups. **(C)** Imaging of Fv/Fm from *Pst* WT- or Δ*hrcC-*infected plants 72 hpi illustrating the TS33 effector-dependent drop on Fv/Fm. Arrows indicate infected leaves. Images are presented in a false color scale (0.85-0.60). For the case of RSH3OX, saturated false blue pixels indicate pixels in which Fv/Fm ≤ 0.6.

### ppGpp is required for chloroplast-mediated resistance against *Botrytis cinerea*

ppGpp-null plants are reported to accumulate higher levels of both SA and JA (Abdelkefi *et al*., 2018; Inazu *et al*., 2024). Since JA signalling is generally protective against necrotrophic pathogens (Glazebrook, 2005), elevated basal JA might be expected to enhance *rshq* resistance to *Botrytis cinerea*; however, given the well-established antagonism between SA and JA pathways (Thaler *et al*., 2012; Pieterse *et al*., 2012), concomitantly elevated SA could instead compromise JA-mediated defence output. Also, we found that RSH3OX and *rshq* plants exhibit differential expression of *SAG13*, *UMAMIT20*, *GRXS13* and *BRG2* (Figure 5A) that are characterized as regulators of defence against *Botrytis* infection (Luo *et al*., 2010; La Camera *et al*., 2011; Dhar *et al*., 2020; Prior *et al*., 2026). We therefore wondered whether ppGpp levels could influence the outcome of infection by this necrotrophic pathogen. To this end, we inoculated Col-0, RSH3OX, and *rshq* plants with *Botrytis cinerea* and quantified lesion area after 72h (Figure 5B-D). RSH3OX plants developed lesions of similar size to Col-0, suggesting that ppGpp over-accumulation does not significantly alter resistance to this pathogen. In contrast, *rshq* plants exhibited significantly larger lesions than Col-0, indicating enhanced susceptibility to *B. cinerea*.

**Figure 5.**
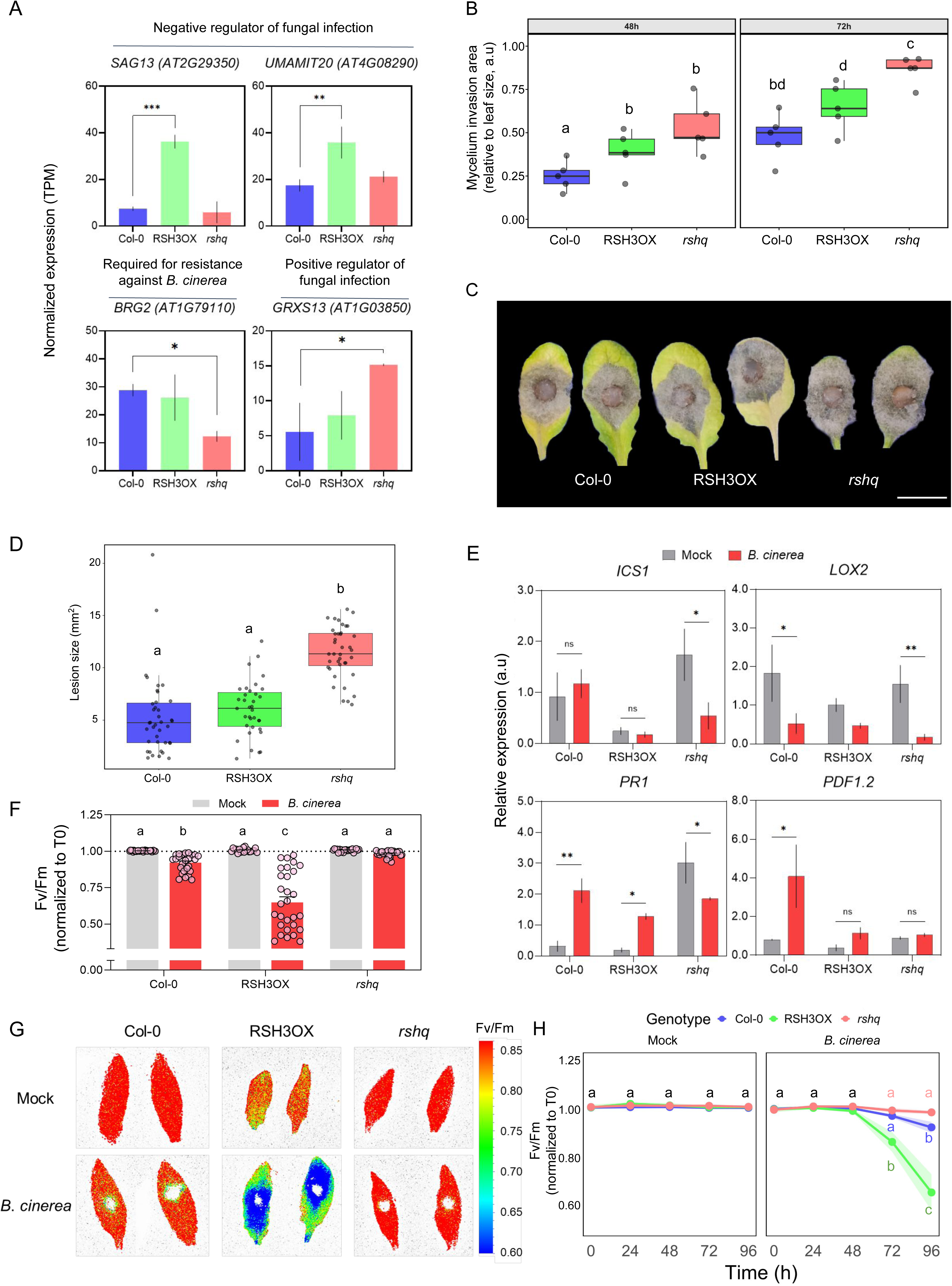
ppGpp levels influence resistance to *Botrytis cinerea*. **(A)** Normalized expression of differentially expressed genes related to fungal infection from RNAseq data. Values are shown in Transcripts per million (TPM). Data is shown in bar plots and presented as mean ± SEM. Asterisks indicate significative differences among genotypes (adj. p-value < 0.05). **(B)** Mycelium invasion area in leaves of Col-0, RSH3OX, and *rshq* plants at 48 and 72 hpi after inoculation with a 5mm agar disc of *Botrytis cinerea* mycelium, expressed relative to total leaf size. Different letters indicate statistically significant differences among genotypes (p < 0.05, one-way ANOVA with post-hoc test). Data are presented in boxplot where the line in the box represents the median and the whiskers represent the 25th and 75th percentile of five biological replicates. **(C)** Representative photographs of detached leaves from Col-0, RSH3OX, and *rshq* plants showing *B. cinerea* mycelium invasion at 72 hpi. Scale bar: 1 cm **(D)** Lesion area in leaves of Col-0, RSH3OX and *rshq* plants at 72 hpi with 10 uL of *B*. *cinerea* spore suspension (10^6^ spores/mL). Different letters indicate statistically significant differences among genotypes (p < 0.05, one-way ANOVA with Fisher post-hoc test). Box plot show the range of distribution of data. The median is represented by black line within the box. **(E)** RT-qPCR analysis of the SA biosynthesis marker *ICS1*, the JA biosynthesis gene *LOX2*, SA-responsive marker *PR1* and JA-responsive marker *PDF1.2* in leaves of Col-0, RSH3OX, and *rshq* plants under mock or *Pst* DC3000 infection conditions. Expression values are shown in arbitrary units (a.u.) relative to *EF1A*. Data are presented as mean ± SEM of biological replicates. **(F)** Photosynthesis efficiency (Fv/Fm) of detached leaves inoculated with Mock (PDB liquid medium) or *B*. *cinerea* spore suspension (10^6^ spores/mL). Data are shown in arbitrary units (a.u.) relative to 0h post-infection (T0). Fv/Fm was measured with a custom mask on FluorCam7 to take into account only pixels around infection site. Data are presented as mean ± SEM. Different letters indicate statistical significant differences among genotypes and treatments (p<0.05, two-way ANOVA with post-hoc test). **(G)** Imaging of Fv/Fm from Mock- or *B. cinerea*-inoculated leaves at 72 hpi illustrating drop on Fv/Fm around the droplet inoculation site. Images are presented in a false color scale (0.85-0.60). For the case of RSH3OX, saturated false blue pixels indicate pixels in which Fv/Fm ≤ 0.6. **(H)** Kinetics of photosynthesis efficiency during *B. cinerea* infection. Data are shown in arbitrary units (a.u.) relative to 0h post-infection (T0). Data are presented in line plots as mean ± SEM. Data from 5F, 5G and 5H are derived from the same experiments.

To determine whether the altered susceptibility to *B. cinerea* was associated with changes in SA- or JA-dependent defence responses, we analyzed the expression of representative marker genes from both pathways (Figure 5E). As expected, we found that upon infection, Col-0 plants showed unaffected expression of the SA biosynthesis marker *ICS1,* whilst expression of the JA biosynthesis marker *LOX2* was suppressed (Windram *et al*., 2012). However, the SA-responsive marker *PR1* and JA-responsive marker *PDF1.2* were both induced in Col-0. This is consistent with reports that SA-associated responses to *B. cinerea* may occur independently of ICS1- mediated SA biosynthesis (Ferrari *et al*., 2003). RSH3OX plants showed a similar transcriptional response to Col-0, consistent with a similar lesion size (Figure 5D). In contrast, *rshq* plants showed higher *ICS1* expression in the mock treatment, and a strong suppression during infection. In line with the increased lesion size (Figure 5D), *PR1* expression was suppressed during infection and, notably, *PDF1.2* was not induced relative to mock, despite the elevated basal JA levels previously reported in ppGpp-null plants (Abdelkefi *et al*., 2018; Inazu *et al*., 2024). This failure to mount a *PDF1.2* response upon infection, rather than basal JA status itself, is consistent with SA-mediated suppression of JA signalling and may account for the enhanced susceptibility of *rshq* plants to *B. cinerea*: while ppGpp depletion in *rshq* plants promotes resistance to the hemi- biotrophic pathogen *Pst* DC3000, it confers greater susceptibility to the necrotrophic pathogen *B. cinerea*, suggesting that ppGpp contributes to the hormonal signalling balance required for broad-spectrum defence competence.

To further investigate the cellular basis of the contrasting defence phenotypes observed across genotypes, we assessed chlorophyll fluorescence at different times post infection. We found that, as for *Pst* infection, Col-0 plants undergo an Fv/Fm decrease around the infection site at 72 hpi compared to the mock treatment (Figure 5F-H). RSH3OX plants exhibited a stronger Fv/Fm decrease around the infection site at 72 hpi. *rshq* plants did not show changes in Fv/Fm bordering the infection site, although an absence of chlorophyll fluorescence due to fungal growth is still observed at the center of the lesion.

These results suggest that ppGpp depletion predisposes plants to abrupt, tightly localized cell death at the infection site, without the progressive spread of photosynthetic dysfunction observed in RSH3OX, potentially reflecting a distinct mode of tissue collapse that may underlie both the restriction of bacterial proliferation and the enhanced susceptibility to *B. cinerea* observed in *rshq* plants.

## Discussion

### ppGpp functions as a chloroplast-based regulator of nuclear defence gene expression

Chloroplast function in plants is rapidly adjusted during pathogen attack. Our findings establish ppGpp as a critical component of chloroplast retrograde signalling that modulates plant immunity by modulating the expression of nuclear-encoded defence genes. The observation that both ppGpp overaccumulation (RSH3OX) and deficiency (*rshq*) trigger widespread transcriptional reprogramming of immunity-related genes demonstrates that precise regulation of ppGpp levels is essential for appropriate defence responses. This is consistent with the emerging view that chloroplasts function not merely as photosynthetic organelles but as metabolic sensors that integrate environmental information and transmit signals to regulate nuclear gene expression (Chan *et al*., 2016; de Souza *et al*., 2017). A related study on the control of the nuclear transcriptome by ppGpp under nitrogen deficiency conditions was published during the preparation of this manuscript, using the same *rsh1-rsh2-rsh3-crsh* quadruple mutant line employed in our own work (*rshq*) (Nemoto *et al*., 2026). The authors found that ppGpp accumulation is required for the transcriptional repression of immune system processes and SA-response genes during nitrogen starvation, a repression that is lost in the ppGpp-null, *rshq* line. This parallels our own findings that elevated ppGpp suppresses SA-associated defence gene induction upon pathogen challenge, while ppGpp deficiency activates these same pathways, reinforcing the view that ppGpp functions as a conserved repressor of SA-linked nuclear transcription across distinct plastid-derived stress signals, and supporting its broader role as a central component of plastid-to-nucleus retrograde signalling in the context of the defence response.

### ppGpp controls SA and JA catabolism

A key finding of this study is that ppGpp regulates the expression of genes encoding hormone- deactivating and activating enzymes. In the case of SA, ppGpp overaccumulation upregulates *BSMT1*, which encodes a methyltransferase that converts SA to the inactive (yet volatile and mobile) form, MeSA (Chen *et al*., 2003). Conversely, ppGpp deficiency upregulates *MES1* and *MES9*, which encode esterases that hydrolyze MeSA back to active SA (Yang *et al*., 2006; Vlot *et al*., 2008). Additionally, low levels of ppGpp downregulate of *UGT74F2*, which encodes a glucosyltransferase that conjugates SA to glucose for storage in the vacuole (Dean & Delaney, 2008). Together, these transcriptional changes shift the equilibrium toward active SA in ppGpp-deficient plants and toward inactive/storage forms in ppGpp-overaccumulating plants, providing a molecular explanation for the altered SA levels previously reported (Abdelkefi *et al*., 2018; Inazu *et al*., 2024).

On the other hand, ppGpp regulates the JA pathway by controlling the expression of genes encoding catabolic JA enzymes. RSH3 overexpression is associated with elevated levels of *CYP94B3*, *JOX2*, *JOX3*, and *JOX4*, which encode enzymes that oxidize JA-Ile to inactive forms (Koo *et al*., 2011; Heitz *et al*., 2012), as well as *ILL6*, which conjugates inactive JA to amino acids, potentially diverting JA away from the bioactive JA-Ile pool (Widemann *et al*., 2013). In contrast, *rshq* plants show evidence of activated JA metabolism, with higher JA levels previously reported (Inazu *et al*., 2024). However, ppGpp deficiency did not significantly alter the expression of any JA catabolic genes (Figure 1D and Table S1), in contrast to its effect specifically upregulating the transcription of genes for SA metabolism (*MES1, MES9, UGT74F2*). This asymmetry, RSH3 overexpression broadly reprogramming JA catabolism at the transcriptional level while ppGpp deficiency specifically upregulates SA metabolism genes without a corresponding transcriptional signature in JA catabolism, suggests that the elevated JA-Ile levels reported in ppGpp-deficient plants (Inazu *et al*., 2024) likely arise through a mechanism other than transcriptional repression of catabolic enzymes. The coordinated regulation of multiple catabolic steps suggests that ppGpp-dependent signaling exerts multilayered control over hormone catabolism in the SA and JA pathways.

### Temporal dynamics of *RSH* genes during pathogen infection

Our kinetic analysis of *RSH* gene expression during *P. syringae* infection reveals that *RSH2* and *RSH3* are rapidly induced, with expression levels rising within 6-24 hours post-infection. This induction occurs in parallel with, or slightly before, the upregulation of SA biosynthesis (*ICS1*) and JA biosynthesis (*LOX2*) genes, suggesting that ppGpp accumulation is an early component of the defence response rather than a downstream consequence of hormone signalling. This temporal pattern is consistent with the hypothesis that pathogen perception triggers chloroplast stress, leading to *RSH2* and *RSH3* expression, ergo ppGpp synthesis, which in turn modulates nuclear defence transcriptome.

The induction of *RSH2/3* by chitin (Supplementary Figure S1A), a pathogen-associated molecular pattern (PAMP), further supports the idea that ppGpp synthesis is integrated into pattern-triggered immunity (PTI). In bacteria, ppGpp accumulation is triggered by diverse stressors including amino acid starvation, oxidative stress, and cell envelope damage (Hauryliuk *et al*., 2015). In plants, pathogen infection causes oxidative bursts, redox imbalances, and metabolic perturbations that could potentially trigger ppGpp synthesis through similar mechanisms (Nomura *et al*., 2012), though this remains to be tested in plants. The fact that *RSH2* is slightly induced in roots by chitin treatment suggests that ppGpp-dependent signalling may also function in root immunity, an area that warrants further investigation (Supplementary Figure S1A).

While these findings support a role for PAMP perception in triggering *RSH2/RSH3* induction, our results also reveal that a fully virulent, T3SS-competent pathogen is required to elicit this response: T3SS-deficient mutants (*ΔhrpA, ΔhrcC*) failed to induce *RSH2/RSH3*, despite still presenting flagellin. This is at first glance difficult to reconcile with the reported sufficiency of flg22 alone to induce *RSH2* within 1 hour (Qiu *et al*., 2023), particularly since the canonical function of T3SS-delivered effectors is to suppress PAMP-triggered immunity. A T3SS-deficient mutant would therefore be expected to fail to suppress PTI rather than fail to elicit it. One possible explanation is that *RSH2/RSH3* induction during live infection requires a more sustained or cumulative PAMP exposure than a single flg22 treatment provides, potentially reflecting differences in bacterial persistence and multiplication between a poorly-proliferating T3SS mutant and a fully virulent strain. Alternatively, effector activity itself, rather than merely its absence being tolerated, may contribute directly to *RSH2/RSH3* induction, suggesting that the transcriptional response to live infection integrates both PAMP- and effector- derived signals, rather than PAMP perception alone.

In addition, we found that *RSH2/RSH3* induction during infection requires SA biosynthesis, as this response is lost in the ICS1-deficient *sid2* mutant (Supplementary Figure S5B). Given that flg22 alone can induce *RSH2/RSH3-*mediated ppGpp production within 1 hour, according to Qiu and colleagues (Qiu *et al*., 2023), likely faster than how SA biosynthesis can be substantially upregulated, this raises the possibility that PAMP-triggered RSH induction and infection-triggered RSH induction proceed through at least partially distinct mechanisms: a rapid, SA-independent route engaged by PAMP perception alone, and a slower, SA biosynthesis-dependent route that predominates over the course of a full infection. Since exogenous SA treatment did not itself induce *RSH2/RSH3* (Supplementary Figure S5A), this requirement for ICS1 more likely reflects a role for the SA biosynthetic pathway, or related metabolites, than canonical SA signalling *per se*. In any case, methodological differences between the treatments used by Qiu *et al*. (2023) and those used here may also contribute to the observed dichotomy.

### Opposing effects of ppGpp on resistance to hemi-biotrophic and necrotrophic pathogens

The functional consequences of altered ppGpp levels are strikingly pathogen-specific. RSH3OX plants exhibit hypersusceptibility to the hemi-biotrophic pathogen *P. syringae*, with increased bacterial growth and severe chlorosis, while *rshq* plants display enhanced resistance to this pathogen. This pattern is broadly consistent with the reduced SA accumulation reported in RSH3OX and elevated SA in *rshq* plants (Abdelkefi *et al*., 2018; Inazu *et al*., 2024). Nevertheless, our data suggest that the mechanistic basis of *rshq* resistance extends beyond canonical SA-dependent immunity. RSH3OX plants show strongly suppressed transcriptional activation of *ICS1, PR1*, *FRK1*, and *LOX2* upon *Pst* inoculation, indicating that ppGpp over-accumulation impairs canonical immune gene induction despite, or perhaps because of, the elevated bacterial loads these plants support. *rshq* plants, by contrast, display intermediate-to-strong induction of these markers, arguing against a straightforward SA-mediated explanation for their enhanced resistance. It has been reported that ppGpp-deficient plants exhibit slightly higher basal SA levels (Abdelkefi *et al.,* 2018). However, in our experimental conditions, *ICS1* steady state abundance in mock-treated *rshq* plants is similar to Col-0, suggesting that transcript abundance alone may not fully capture differences in SA accumulation or pathway activity between genotypes. Given that SA signalling promotes programmed cell death (Radojcic *et al*., 2018), we hypothesize that an early and localized cell death response to *Pst* might be the primary mechanism restricting bacterial proliferation in *rshq* plants. We did not directly assess cell death in this context, and this remains speculative (Wolfgang *et al*., 2011). Nonetheless, a related possibility emerges from our *B. cinerea* infection data: *rshq* plants show abrupt, tightly localized photosynthetic collapse at the infection site without progressive spread to surrounding tissue, in contrast to the more gradual, spreading dysfunction observed in RSH3OX plants. If ppGpp deficiency indeed predisposes plant tissue to a faster or more contained mode of cell death upon pathogen challenge, this could plausibly explain two superficially contrasting phenotypes at once: restricting the hemi-biotrophic pathogen *Pst*, which depends on living host tissue in its first phase, while increasing susceptibility to the necrotrophic pathogen *B. cinerea*, which thrives on dead tissue. Under this model, ppGpp would not function as a simple positive or negative regulator of defence, but rather as a determinant of the mode-of-tissue response to infection, with opposite consequences depending on pathogen lifestyle.

Although impaired callose deposition was observed in both RSH3OX and *rshq* plants, and therefore cannot alone account for their opposing susceptibility phenotypes, it nonetheless indicates that high ppGpp levels compromise this branch of cell wall-based defense, potentially through effects on sugar metabolism or vesicle trafficking (Luna *et al*., 2011). In contrast, the attenuated *Pst*-induced stomatal closure, along with a constitutively wider stomatal aperture, was specific to RSH3OX plants, strongly suggesting that ppGpp levels negatively regulate stomatal closure, conceivably due to *BSMT1* upregulation and the downregulation of SA signalling and *FRK1* (Zheng *et al*., 2012). Taken together, these results suggest that RSH3OX hypersusceptibility to *Pseudomonas* infection stems primarily from this genotype-specific impairment in stomatal immunity, compounding the broader suppression of PTI-associated gene induction, rather than from the callose defect shared with *rshq*. However, *rshq* resistance to *Pst* cannot be merely explained by an exacerbated PTI response. In addition, *Pst* specifically upregulates expression of *PDF1.2* in *rshq* plants. Since *PDF1.2* encodes a cysteine-rich antimicrobial peptide, its early induction may also contribute to restricting bacterial proliferation in *rshq* plants, alongside the cell death-biased response proposed above.

In contrast, *rshq* plants are hypersusceptible to *Botrytis cinerea*, a necrotrophic fungus. Strikingly, *rshq* plants show suppressed transcriptional induction of *PR1* and, notably, *PDF1.2* during fungal infection, the latter representing a clean reversal of the strong *PDF1.2* induction observed in *rshq* upon *Pst* infection (Figure 3C and 5E). *PDF1.2* is a well-established antifungal defensin systemically activated by fungal pathogens, including *B. cinerea* (Manners *et al.,* 1998; Thomma *et al*., 1998). This suggests that the failure to induce *PDF1.2*, specifically upon fungal challenge, despite its robust induction during bacterial infection, may directly contribute to the hypersusceptibility of *rshq* plants to *B. cinerea*.

Our results are consistent with previous reports on the impact of viral infection on RSH3OX and ppGpp-null plants (Abdelkefi *et al.,* 2018). While RSH3OX plants exhibit hypersusceptibility to TuMV, ppGpp-null plants restrict virus accumulation. This parallel reinforces our model, given that viruses, like the biotrophic phase of hemi-biotrophic pathogens, depend entirely on living host tissue for replication.

The upregulation of *SAG13* in RSH3OX plants is also noteworthy in this context. SAG13 has been reported to negatively regulate bacterial resistance while positively regulating fungal resistance (Seltmann *et al*., 2010). While *SAG13* upregulation is consistent with the *Pst* hypersusceptibility phenotype, RSH3OX plants do not display enhanced resistance to *B. cinerea* despite elevated *SAG13* expression. This indicates that the anti-fungal effect of SAG13 may be overridden by other ppGpp-dependent changes in RSH3OX plants during infection, and that ppGpp modulates defence outputs through different mechanisms.

Taken together, these findings highlight a critical role for ppGpp in tuning the balance between hormone-mediated transcriptional immunity and cell death-based resistance. Rather than acting as a simple regulator of the SA-JA antagonism, ppGpp appears to influence the fundamental mode of immune execution: determining whether plants respond to pathogens through controlled transcriptional reprogramming or through a broader response with cell death as its endpoint. The conditions that trigger ppGpp accumulation in nature, such as light stress and nutrient limitation, may therefore have profound consequences not only for growth-defence trade-offs but for the qualitative nature of the immune response itself.

### ppGpp as a metabolic checkpoint linking chloroplast status to immunity

The integration of ppGpp signalling with defence responses may reflect the fundamental dependence of plant immunity on chloroplast-derived metabolites and energy. Effective immune responses require massive metabolic reprogramming, including the production of antimicrobial compounds, cell wall reinforcements, and reactive oxygen species (Bolton, 2009; Berger *et al*., 2007). These processes are energetically expensive and draw heavily on photosynthetically-derived ATP, NADPH, and carbon skeletons. ppGpp, as a chloroplast-localized signal of metabolic status, may function as a checkpoint that modulates the activation and mode of defence programs in accordance with chloroplast functional state.

ppGpp is known to protect photosystem II and promote resistance to high light stress (Romand *et al*., 2025). In this work we extended this role to pathogen infection: both *Pst* and *B. cinerea* induce a drop in photosynthesis efficiency, and ppGpp status directly determines the magnitude and localization of this decline. During bacterial infection, photosynthesis efficiency is regulated by T3SS-dependent virulence activity. PAMPs alone are not sufficient, as *RSH2* and *RSH3* expression is not induced by *ΔhrcC* infection and Fv/Fm does not change during either *ΔhrcC* infection or flg22 infiltration (Figure 4B and Supplementary Figure S8). RSH3OX plants show an earlier and more severe Fv/Fm decline than Col-0 under *Pst* infection, while *rshq* plants show no decline at all (Figure 4A), indicating that ppGpp is required for this pathogen-triggered photosynthetic collapse. A similar, genotype-dependent pattern emerges during *B. cinerea* infection: RSH3OX plants show an intensified, spreading loss of photosynthetic efficiency around the infection site, whereas *rshq* plants show no decline at the lesion border at all, despite being the more susceptible genotype to this pathogen (Figure 5). We can solve this apparent paradox with the cell death-biased model proposed above: rather than a gradual, spreading photosynthetic decline, *rshq* tissue may undergo abrupt, tightly localized collapse that does not propagate outward, distinguishing it mechanistically from the progressive dysfunction seen in RSH3OX. Notably, RSH3 specifically, rather than RSH2, is the paralog transcriptionally induced by *B. cinerea* infection (Figure 2C), and RSH3OX is also the genotype exhibiting the most pronounced local photosynthetic collapse under this pathogen, suggesting a direct link between which RSH paralog is engaged by a given pathogen and the resulting chloroplast-level phenotype.

To test whether ppGpp acts as a direct, causal determinant of infection outcome we used a dexamethasone-inducible line expressing a chloroplast-targeted bacterial ppGpp synthase, the SYN line (Sugliani *et al*., 2016), to acutely elevate ppGpp levels in an otherwise wild-type background. Acute ppGpp induction alone reduced plant growth, but its combination with *Pst* infection produced a significantly greater reduction in plant size, demonstrating that an acute increase in ppGpp is sufficient to exacerbate pathogen-induced damage independent of any long-term developmental consequences of chronically altered ppGpp levels. This supports a direct, mechanistic role for ppGpp in shaping infection outcome, consistent with its proposed function as a checkpoint linking chloroplast status to immune execution. Together with the disruption *PDF1.2* induction observed in ppGpp-altered lines (Figures 3D and 5E; Nemoto *et al.,* 2026), these findings suggest that ppGpp does not simply gate the activation of defence programs but influences the fundamental mode of immune execution: high ppGpp levels suppress transcriptional defence activation, rendering plants unable to mount effective canonical immune responses, while ppGpp depletion permits partial transcriptional immunity against *Pst* and susceptibility to *Botrytis*.

Interestingly, our transcriptomic data reveal that ppGpp regulates not only hormone metabolism but also genes involved in terpenoid biosynthesis, wounding responses, and ethylene signalling, all processes with known roles in defence. Terpenoids include a diverse array of antimicrobial compounds and signalling molecules (Tholl, 2015), while ethylene cooperates with JA in defence against necrotrophs and herbivores (Pieterse *et al*., 2012). The broad scope of ppGpp-responsive genes suggests that this molecule functions as a wide-ranging regulator that coordinates multiple defensive and metabolic pathways in response to chloroplast status, with consequences that extend well beyond the SA-JA hormonal balance.

### Evolutionary perspective: ppGpp as a conserved stress signal

RSH genes are thought to have entered the proto-plant lineage not through vertical inheritance from the cyanobacterial endosymbiont that gave rise to chloroplasts, but through at least one, and likely several, independent horizontal gene transfer events from different bacterial *phyla* (Atkinson *et al*., 2011; Ito *et al*., 2017; Avilan *et al*., 2019). In bacteria, ppGpp coordinates the transition from exponential growth to survival mode under nutrient limitation, balancing ribosome biogenesis, amino acid synthesis, and stress resistance (Hauryliuk *et al*., 2015). The repurposing of this laterally acquired bacterial signalling system in plants to regulate nuclear gene expression and immune outcomes represents a remarkable example of cross-kingdom gene transfer being co-opted into the complex regulatory networks governing multicellular immunity.

Key differences nevertheless exist between bacterial and plant ppGpp signalling. In bacteria, ppGpp directly binds RNA polymerase to regulate transcription (Ross *et al*., 2013), whereas in plants, ppGpp regulates plastid gene expression through mechanisms that remain incompletely understood and appears to trigger retrograde signals that regulate nuclear transcription indirectly. The molecular nature of these retrograde signals remains a critical open question. Potential mechanisms include ppGpp-regulated metabolites such as isoprenoids or tetrapyrroles, reactive oxygen species generated downstream of altered chloroplast metabolism, or changes in redox status (Chan *et al*., 2016; Larkin, 2016).

### Unanswered questions and future directions

Several key questions emerge from this work. First, what is the molecular mechanism by which chloroplast-localized ppGpp influences nuclear gene expression? Does ppGpp itself exit the chloroplast, or does it trigger the production of secondary signals? Candidate retrograde signals include metabolites such as methylerythritol cyclodiphosphate (MEcPP), β-cyclocitral, or changes in chloroplast redox status (Xiao *et al*., 2012; D’Alessandro *et al*., 2018). Testing whether ppGpp regulates the levels of known retrograde signals would help position ppGpp within the broader network of organellar-nuclear communication.

Second, how do different stresses like pathogen infection, light transitions, and nutrient limitation, regulate ppGpp synthesis? In bacteria, ppGpp is synthesized in response to, for example, uncharged tRNAs accumulating at the ribosome (Hauryliuk *et al*., 2015). Whether a similar mechanism operates in plant chloroplasts, or whether ppGpp synthesis responds to other signals such as ATP/ADP ratios, redox status, or specific metabolites, remains unclear.

Third, does ppGpp regulate immunity in other plant species and in agricultural contexts? If ppGpp functions as a general regulator of stress-defence trade-offs, manipulating ppGpp levels could potentially be used to modulate disease resistance in crops. However, the opposing effects on biotrophic and necrotrophic resistance observed here, and the growth defects associated with ppGpp over-accumulation (Sugliani *et al*., 2016), suggest that any such applications would require careful consideration of pathogen context and would likely necessitate inducible or tissue-specific expression systems or the identification and engineering of the specific regulatory mechanisms.

## Conclusion

This study establishes ppGpp as a critical regulator of plant immunity that operates by controlling the expression of nuclear-encoded genes involved in plant immunity pathways. Our data reveal that ppGpp does not simply tune the magnitude of immune responses but influences their fundamental mode of execution: elevated ppGpp levels suppress transcriptional defence outputs and compromise effective pathogen restriction, whereas ppGpp depletion suppresses canonical hormone-mediated immunity. The opposing susceptibility phenotypes of RSH3OX and *rshq* plants to biotrophic and necrotrophic pathogens further underscore the importance of ppGpp in matching defence responses to the nature of the pathogen threat. Together, our findings support a model in which chloroplast-localised ppGpp serves as a metabolic checkpoint that integrates information about organellar status with nuclear defence programmes, with consequences that extend beyond the SA-JA hormonal balance. Furthermore, ppGpp directly regulates the plant’s immune response, as an acute increase in ppGpp levels is sufficient to exacerbate pathogen-induced damage. Understanding the molecular mechanisms linking ppGpp to retrograde signalling and defence gene regulation will be critical for developing strategies to enhance crop immunity while maintaining photosynthetic efficiency and growth.

## Materials and methodology

### Plant material and growth conditions

The *Arabidopsis thaliana* ecotype Columbia-0 (Col-0) was used as wild type (WT) in all the experiments. 35S::RSH3-GFP (RSH3OX) and DEX-inducible SYN plants (pOPOn2.1::RBCS1A^1-80^-relA^1-386^) were reported in Sugliani *et al.,* (2016) and the *rsh1rsh2rsh3crsh* quadruple mutant line (*rshq)* was kindly provided by Dr. Shinji Masuda (Inazu *et al.,* 2024). *sid2* and *jar1* mutant lines were kindly provided by Dr. Gustavo Gudesblat (iB3, FCEN-UBA). Plants were grown in long day conditions (16:8 light/dark) at 21°C and light intensity of 100-120 μmol s^−1^ m^−2^. Adult plants used in experiments were germinated and grown in soil for 3-4 weeks. For in-plate experiments, seeds were sterilized with bleach 30% v/v during 5 minutes, rinsed with sterilized water and sown in half-strength Murashige and Skoog (MS) medium supplemented with 2-(N-morpholino)ethanesulfonic acid (MES) 0.03% w/v containing 1.5% w/v agar.

### Total RNA isolation and mRNA level quantification

Total RNA was isolated from adult plant leaves or whole seedlings. Briefly, plant material was flash frozen in liquid nitrogen and stored at -80°C. RNA was extracted using TRIzol, integrity was evaluated through agarose electrophoresis and quantification was performed with a Nanodrop 1000 spectrophotometer (Thermo scientific). cDNA synthesis of polyadenylated transcripts was performed using 500 ng of total RNA and Moloney Murine Leukemia Virus Retrotranscriptase (MMLV-RT) according to manufacturer specifications (Invitrogen). Transcript levels were assessed by quantitative PCR of cDNA (RT-qPCR) according to manufacturer specifications (FastStart® Universal SYBR® Green Master(Rox), Roche) in a Eppendorf RealPlex 2 Real time PCR machine. Primers used in this study are listed in Table S2.

### Plant material for RNA sequencing (RNA seq)

Seeds were sown on MS-MES agar medium and stratified for 3 days at 4°C. 14-day-old seedlings grown in continuous light were incubated in the dark for 48h. Then, they were transferred to light (light) or maintained in dark (dark) for an additional 4h. RNA was extracted using Spectrum^TM^ Plant Total RNA Kit (Sigma Aldrich) following manufacturer specifications. 150 pb paired-end sequencing was carried out in a Novaseq Illumina sequencer (HWI-ST1276, Novogene) yielding ∼40 million reads per sample.

### RNAseq data processing

Raw reads quality was assessed with FastQC. Subsequently, reads were filtered (<50pb) and adapters were trimmed using TrimGalore (https://github.com/FelixKrueger/TrimGalore). Clean reads were aligned to the TAIR10 reference genome using STAR aligner with options `-- runThreadN 14 --alignSJDBoverhangMin 1 --alignIntronMax 5000 -- outSAMtype BAM SortedByCoordinate --outWigType wiggle -- outWigStrand Unstranded –-quantMode GeneCounts`. Reads were quantified and normalized using salmon software (Patro *et al*., 2017) against the AtRTD2_QUASI reference transcriptome (Zhang *et al*., 2017). Salmon output files were used as input in 3D RNA Seq App (Guo *et al*., 2020) for differential gene expression analysis. Read counts and counts per million (CPM) were obtained with TXIMPORT (Bioconductor) in R and the lengthScaledTPM method (Soneson, *et al.,* 2016). Low expressed transcripts were filtered using the mean-trend variance method (CPM ≥ 2 in at least three samples) and differential gene expression was considered for genes with |Log_2_(Fold Change)| ≥ 1. For statistical significance, a Benjamini & Hochberg approach was used for multiple-testing correction. False Discovery Rate (FDR) < 0.05 was considered for differential gene expression analysis. Differentially expressed genes (DEG) from our RNA seq data can be found in Table S1.

### Functional enrichment and gene ontology (GO) analysis

Functional enrichment of RNA seq data was evaluated using PantherDB Classification System (Mi *et al.,* 2019). The bubble plot of GO term enrichment was constructed with a custom Python script using the seaborn library.

### Public RNAseq data analysis

Publicly available RNAseq data were retrieved from SRA with `sra-toolkit` (prefetch `-- option-file`) and sra files were converted to fastq files with `fasterq-dump`. Read quality assessing, adapter trimming and downstream analysis were performed as described above.

### Microarray data analysis

Microarray data (GSE56094) were retrieved from Gene Expression Omnibus (GEO). Briefly, a series matrix file from data was downloaded and Complete Arabidopsis Transcriptome Microarray Data (CATMAs) IDs were assigned to each gene locus (TAIR9). Then, Lowess normalized signal data (Berger, *et al.,* 2004) were normalized to T0 (0 hpi).

### Single cell RNAseq(sc-RNA seq) analysis

sc-RNA seq data from *Arabidopsis thaliana* plants infected with *Pseudomonas syringae* DC3000 were retrieved from GEO (GSE213622; Zhu *et al.,* 2023). Data were processed using Seurat v5.3.1 in R. Cell type identity was assigned using the metadata column `predicted.id`. Additionally, Seurat clusters were annotated according to the original article nomenclature (M1– M18, B13, P14, G15, C16). Dimensionality reduction and visualization were performed using UMAP embeddings pre-computed in the original dataset.

Gene expression data were normalized using a reads per cell scaling approach (scale factor = 10000; Transcripts per ten thousand, TP10K) to preserve expression values in a linear scale.

Differential expression analysis between DC3000- and mock-inoculated plant cells was performed for each cell type using Wilcoxon test implemented in FindMarkers (Seurat) with a minimum cell fraction threshold of 10% (`min.pct=0.1`) and no log-fold change pre-filter (`logfc.threshold=0)`. Results were corrected for multiple testing using Bonferroni method (default for Seurat). Genes with an adj. P-value < 0.05 were considered differentially expressed. Log2FC were visualized as heatmaps. For this article, we focused on the expression of *RSH1, RSH2, RSH3* and *cRSH*.

### Pseudomonas syringae infection

*Pseudomonas syringae* pvar. *tomato* DC3000 (wild type) and Δ*hrcC* strain were used for pathogen infection assay to evaluate bacterial growth. Bacteria were grown in Luria-Bertani (LB) agar 1,5% supplemented with kanamycin 75 μg/mL and rifampicin 100 μg/mL for 48h at 28°C. Bacteria were infiltrated with a needless syringe and MgCl_2_ 10mM was used as a mock. For bacterial growth *in planta*, 7mm discs were perforated from adult (3-to-4 week old) plants and homogenised in MgCl_2_ 10 mM with our tissue disruptor (Gonzalez *et al*., 2026). Bacterial growth was then assessed by counting colony forming units (CFU/mL) by plate dilution. For RNA experiments, mock- or pathogen-infiltrated leaves were harvested and flash frozen in liquid nitrogen at 24 hpi.

### Callose quantification

Callose deposition was evaluated by infecting adult plants with a Type-III-secretion system (T3SS)-deficient strain (*ΔhrcC*) of *P*. *syringae* DC3000. Briefly, plants were infiltrated with bacteria at OD_600_=0.1 and 24 hours post-infection, leaves were harvested in ethanol for chlorophyll removal. Then, leaves were submerged in a callose staining buffer (aniline blue 0,01% (w/v) and K_2_HPO_4_ 150 mM). Callose deposition was evaluated from leaves of nine different plants for each genotype. Images were taken with an Axioplan epifluorescence microscope (Zeiss, Germany) from Universidad Nacional de Córdoba, Argentina and deposits were quantified with Fiji-based software CalloseMeasurer (Zhou *et al*., 2012). Results are expressed as the number of callose deposits relative to untreated plants.

### Stomatal aperture assay

Stomatal aperture assays were performed according to Pantaleno *et al*., (2024). Epidermal peels from abaxial side of 5- or 6-week-old plants were excised with tweezers and incubated in opening buffer (MES 5 mM pH=6.1 and KCl 50 mM) for 3h under light (200 *μ*mol m^-2^ s^-1^) and maintained in the same buffer (mock) or exposed to *Pst* (OD_600_= 0.1). Stomata were photographed with an AmScope MU1000 camera coupled to an Olympus CKX53 microscope with a 40X lens (LUCPlanFLN, 0.6 numerical aperture). Stomatal aperture width was measured as the maximal distance between the inner walls of guard cells. Measurement was done using ImageJ analysis software (NIH, Bethesda, MD, USA).

### Botrytis cinerea infection

*Botrytis cinerea* infection for phenotyping was performed using the agar disc method. Briefly, *B. cinerea* was grown in Potato Dextrose Agar (PDA) for 14 days at 20 ± 1 °C. Prior to infection, fungi plates were exposed to 4 °C during 4 h for spore release. Then, a 1-mm-agar disc was placed in detached leaves. The lesion area was assessed through mycelium invasion. For experiments with RNA extraction and Chlorophyll fluorescence measurement, detached leaves were inoculated with a 10 μL droplet of *B. cinerea* (10^6^ spores/mL in liquid PD medium). For RNA experiments, mock or infected leaves were harvested and flash frozen in liquid nitrogen at 24 hpi.

### Chlorophyll fluorescence measurement

Plants were transferred to dark for 20 min (center reaction closing) and chlorophyll fluorescence was measured in a Fluorcam FC 800-O imaging fluorometer (Photon System Instruments). PSII maximum quantum yield (Fv/Fm) was calculated as (Fm − F0)/Fm. For the case of *Pst* infected plants, only infected leaves were measured. In the case of *B*. *cinerea* infected leaves, a Fluorcam7-made custom mask was used to measure Fv/Fm around the infection site.

### Flagellin 22 (flg22) infiltration assay

Flg22 (QRLSTGSRINSALDDAAGLQIA, Tebu-Bio, France) was used to monitor Fv/Fm in absence of other bacterial PAMPs and effectors. Briefly, adult plants were infiltrated with flg22 1 *μ*M or H_2_O as mock in the abaxial surface of the leaves.

### Statistical analyses

Statistical analyses were performed with InfoStat (DiRienzo *et al.,* 2018) (http://www.infostat.com.ar).

## Data availability

Raw RNA seq data generated in this study are deposited under the following accession number: PRJNA1496939.

Public RNA seq data used and re-analyzed in this study is deposited under the following accession numbers: PRJNA1253890, PRJNA1158549, GSE56094, GSE213622.

## Gene IDs

Full name and IDs for genes assessed in this study are listed in Table S3..

## Acknowledgements

The authors would like to thank all Argentinian colleagues around the world that contributed to this work either by providing reagents, helpful discussions or even sharing paywalled papers. Moreover, the authors would like to thank MSc. Juan Pablo Fabroni, MSc. Edgar Martinez Moyano and Dr. Rita Ulloa for their assistance with *Botrytis cinerea* experiments. This work was supported by a grant from the International Centre for Genetic Engineering and Biotechnology (ICGEB CRP/ARG22-03 to EP) and funding from the Agence National de la Recherche (ANR-22- CE20-0033). Unfortunately, national science funding for research projects in Argentina has been completely abolished by the national government. As an example, our ANPCYT PICT-2019- 01690 grant was cancelled before the funds were available to us. Due to this, some parts of this work were self-funded by the authors. FEA, ILSJ, FSR and PJS are CONICET fellows. Also, FEA was awarded a UBAINT fellowship to do part of this work in LGBP-CNRS/AMU, Marseille, France. BF is a CNRS researcher. DS, CGM, RST, NMC and EP are CONICET researchers.

## Author contributions

EP, NMC, BF and FEA conceived and designed the experiments. FEA and EP performed the experiment for RNAseq. FEA, PJS and EP performed the bioinformatic analyses. FEA, IJSL, FSR, RST, CB and NMC performed pathogen infection experiments. DS and CGM performed stomata experiments. FEA and BF performed chlorophyll fluorescence experiments. FEA, FSR and RST carried out the RT-qPCR experiments. FEA, NMC, BF and EP interpreted the data. FEA prepared the figures. NMC, BF and EP got the funding. FEA and EP wrote the manuscript with major contributions from NMC and BF and minor contributions from the rest of the authors. All the authors read and reviewed the final manuscript.

---

**Supplementary Figure S1.**
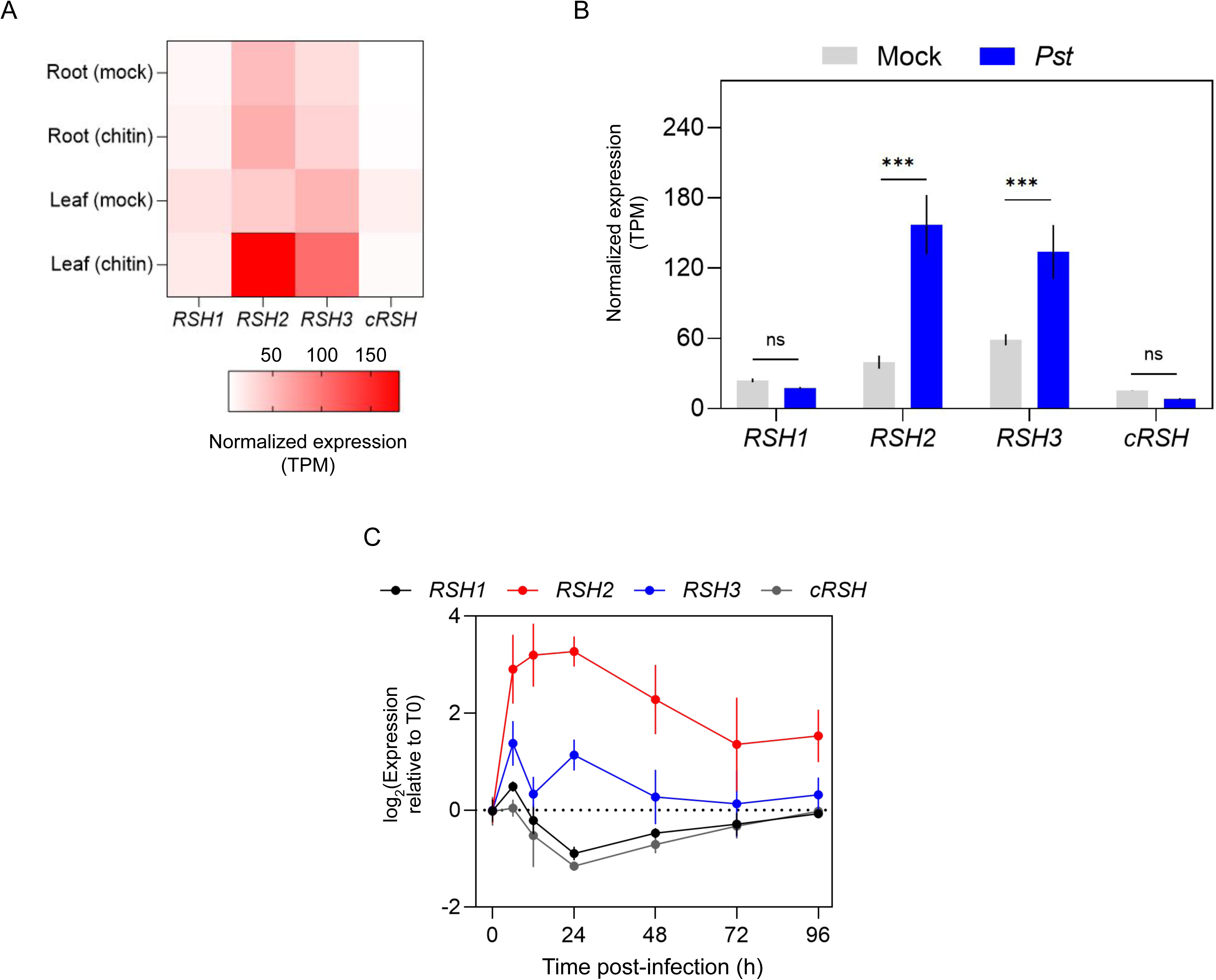
Transcriptomic analysis from public RNAseq data of *RSH1/2/3* and *cRSH* gene expression from *Arabidopsis thaliana* adult plants in immunity-related contexts. **A)** Heatmap of normalized expression of *RSH1*, *RSH2*, *RSH3*, and *cRSH* in roots and leaves of *A. thaliana* plants treated with mock or chitin as indicated. Data are derived from Mackechemu, *et al*. (2025). **B)** Bar plot of normalized expression of *RSH1, RSH2*, *RSH3* and *cRSH* in leaves of adult *A. thaliana* plants inoculated with *Pst* compared to mock-inoculated controls. Values are shown in Transcripts per million (TPM). Data is shown in bar plots and presented as mean ± SD. Asterisks indicate adjusted p-value < 0.05. Data are derived from Mackechemu, *et al*. (2025). **C)** Normalized expression (relative to T0 post-infection) of *RSH1, RSH2, RSH3* and *cRSH* during infection *Botrytis cinerea* infection. Data are derived from public transcriptomic data (Wei, *et al.,* 2024).

**Supplementary Figure S2.**
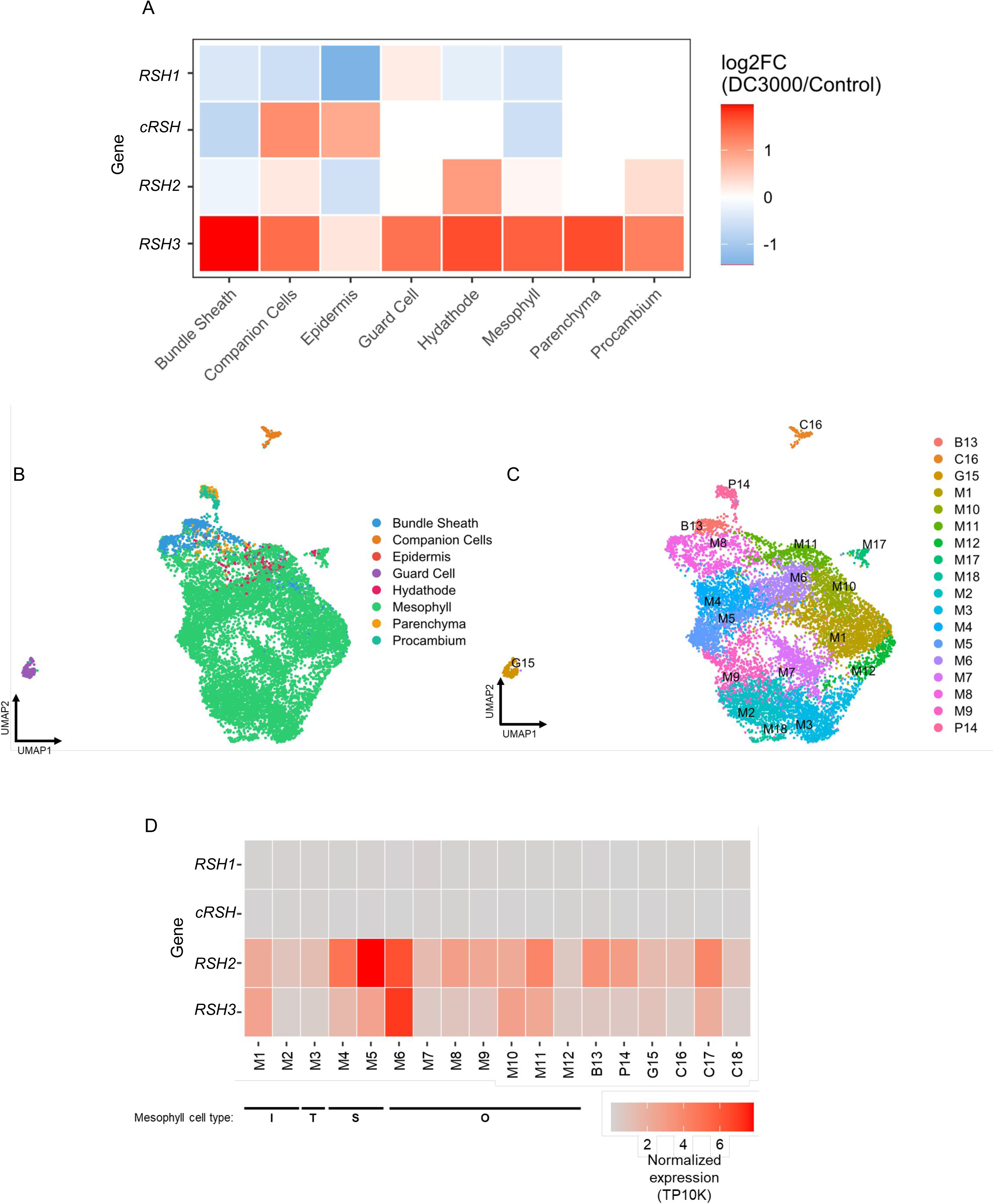
Transcriptomic analysis of public sc-RNAseq data from *Arabidopsis thaliana* adult leaves infected with *Pst*. Uniform Manifold Approximation and Projection (UMAP) from sc-RNAseq (GSE226826, Zhu, *et al*. 2023) colored in base of (A) Cell Type and (B) Cluster identity defined by Zhu, *et al.,* (2023). B: Bundle sheath; C: Companion cells; G: Guard Cells; M: Mesophyll; P: Parenchyma and Procambium. (C) Heatmap plot of differential expression of *RSH1*, *RSH2*, *RSH3*, *cRSH* across all cell types from figure A. (D) Heatmap plot of normalized expression of *RSH1, RSH2, RSH3* and *cRSH* in cluster-separated cell types during pathogen response (sc-RNAseq). Mesophyll cell type are indicated with black line: I: Immunity cell type; T: Transition cell type; S: susceptibility cell type; O: other. Data are presented as TP10K (Transcript per ten thousand transcripts). Data are derived from publicly available transcriptomic dataset: Zhu, *et al*. (2023): GSE226826.

**Supplementary Figure S3.**
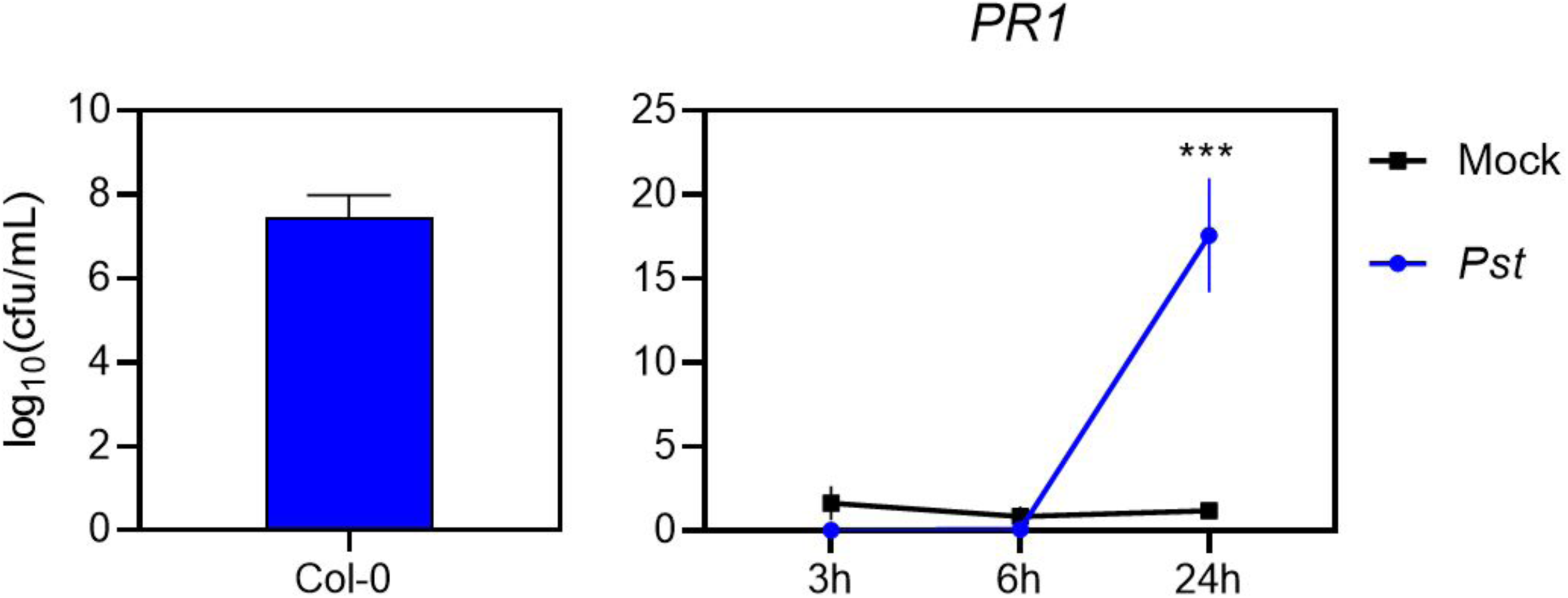
Bacterial growth and plant transcriptional responses during the onset of *Pst* infection. Left panel: Bacterial growth assessment at 3 dpi from time-course experiment (Fig. 2B). Data are presented as mean ± SEM. Right panel: *PR1* transcript levels in *A. thaliana* leaves at 3, 6, and 24 hours after inoculation with *Pst* (blue, OD_600_= 0.0005) or mock solution (black). Expression values are shown in arbitrary units (a.u.) relative to reference gene *EF-1A*. Data are presented as mean ± SEM of four biological replicates. Asterisks indicate statistically significant differences between treatments within each genotype (p < 0.05, two way ANOVA with post-hoc test).

**Supplementary Figure S4.**
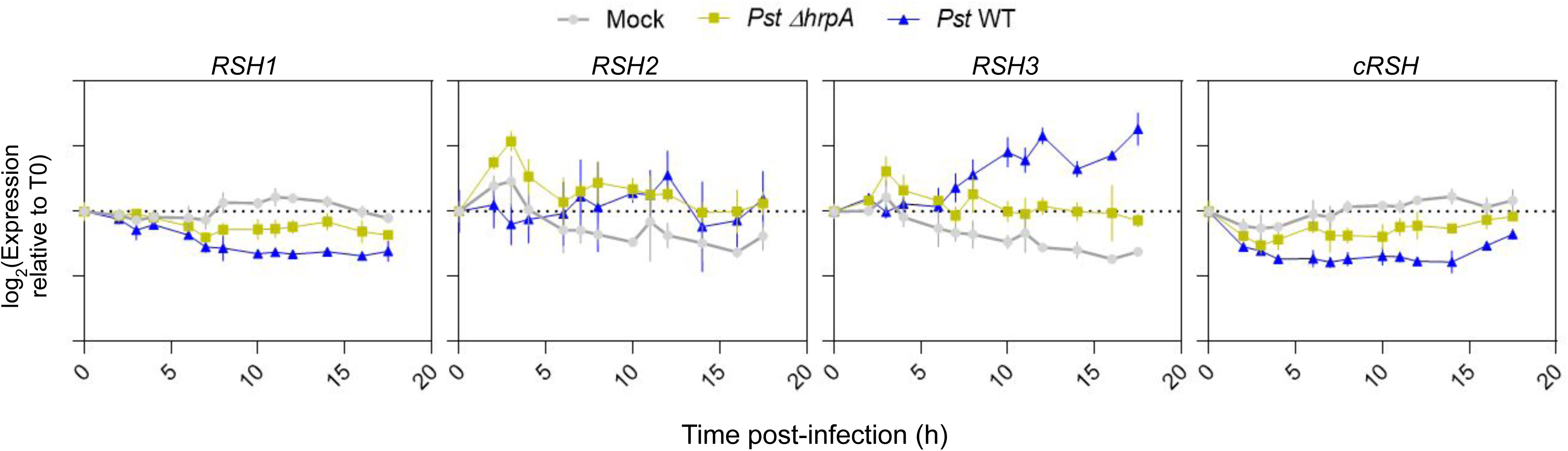
Transcriptomic analysis of public microarray data from *Arabidopsis thaliana* adult leaves infected with *Pst* WT and Δ*hrpA.* Expression dynamics of *RSH1*, *RSH2*, *RSH3*, and *cRSH* over a time course of 17.5 hours post-inoculation with mock (grey), the type III secretion-deficient non-pathogenic *Pst* mutant Δ*hrpA* (yellow), or virulent DC3000 (blue). Values are shown as log₂ expression relative to time zero (T0). Data are presented in line plots as mean ± SD of biological replicates. Data are derived from publicly available transcriptomic datasets (Lewis, *et al*. (2015): GSE56094).

**Supplementary Figure S5.**
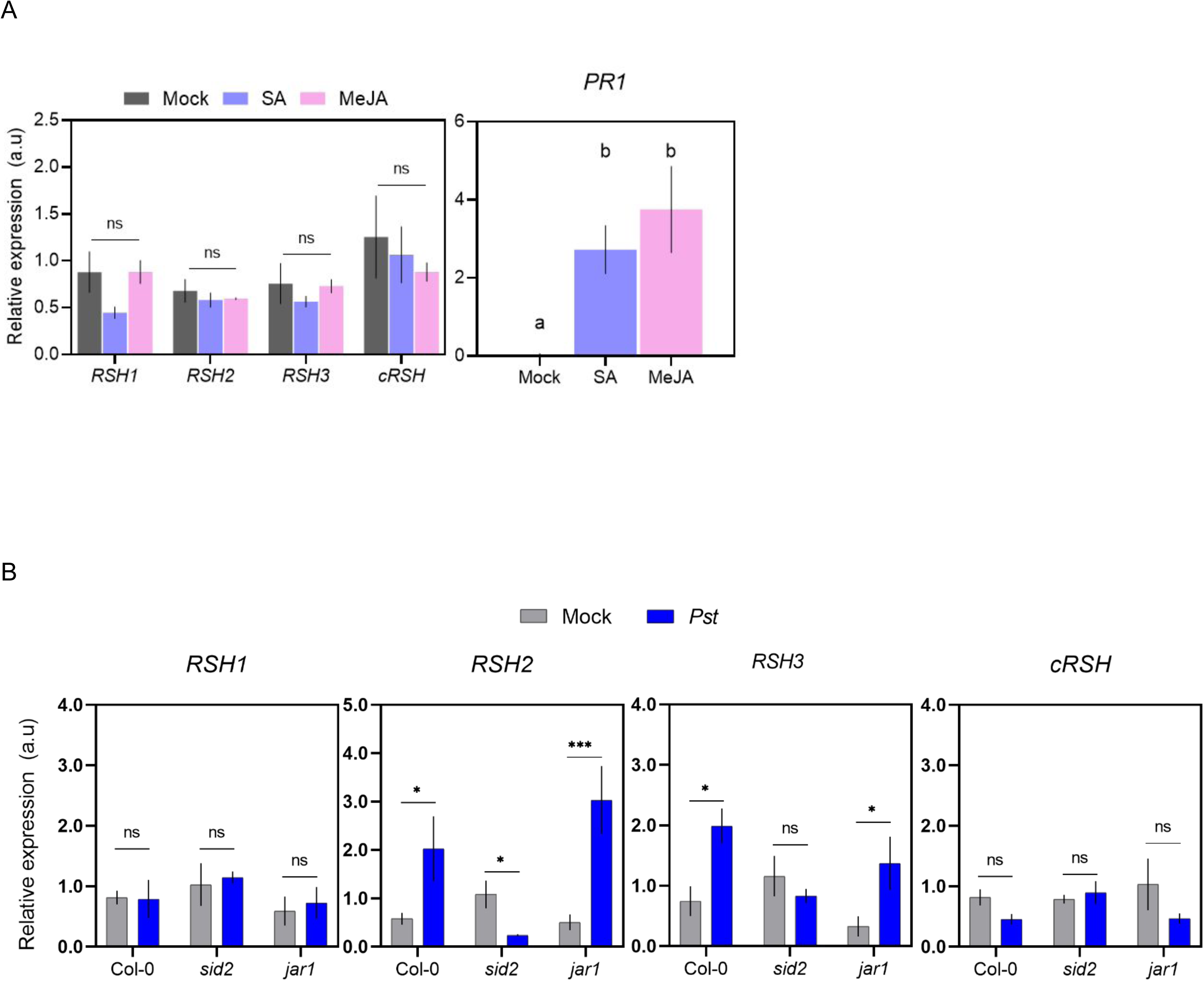
Salicylic acid is necessary but not enough to induce *RSH2* and *RSH3* expression. **(A)** Left panel: RT-qPCR analysis of *RSH1, RSH2, RSH3* and *cRSH* transcript levels in 14-day-old seedlings treated with SA (violet) or MeJA (pink). Right Panel: RT-qPCR analysis of *PR1* as positive control of the treatment. Expression values are shown in arbitrary units (a.u) relative to *EF1-A*. Data are presented as mean ± SEM of three biological replicates. **(B)** RT-qPCR analysis of *RSH1*, *RSH2*, *RSH3* and *cRSH* transcript levels in *A. thaliana* adult plants infiltrated with Mock (gray) or *Pst* (blue) (OD_600_=0.0001) in Col-0, SA-deficient line *sid2* and JA-Ile-deficient line *jar1*. Expression values are shown in arbitrary units (a.u.) relative to *EF1-A*. Data are presented as mean ± SEM of three biological replicates.

**Supplementary Figure S6.**
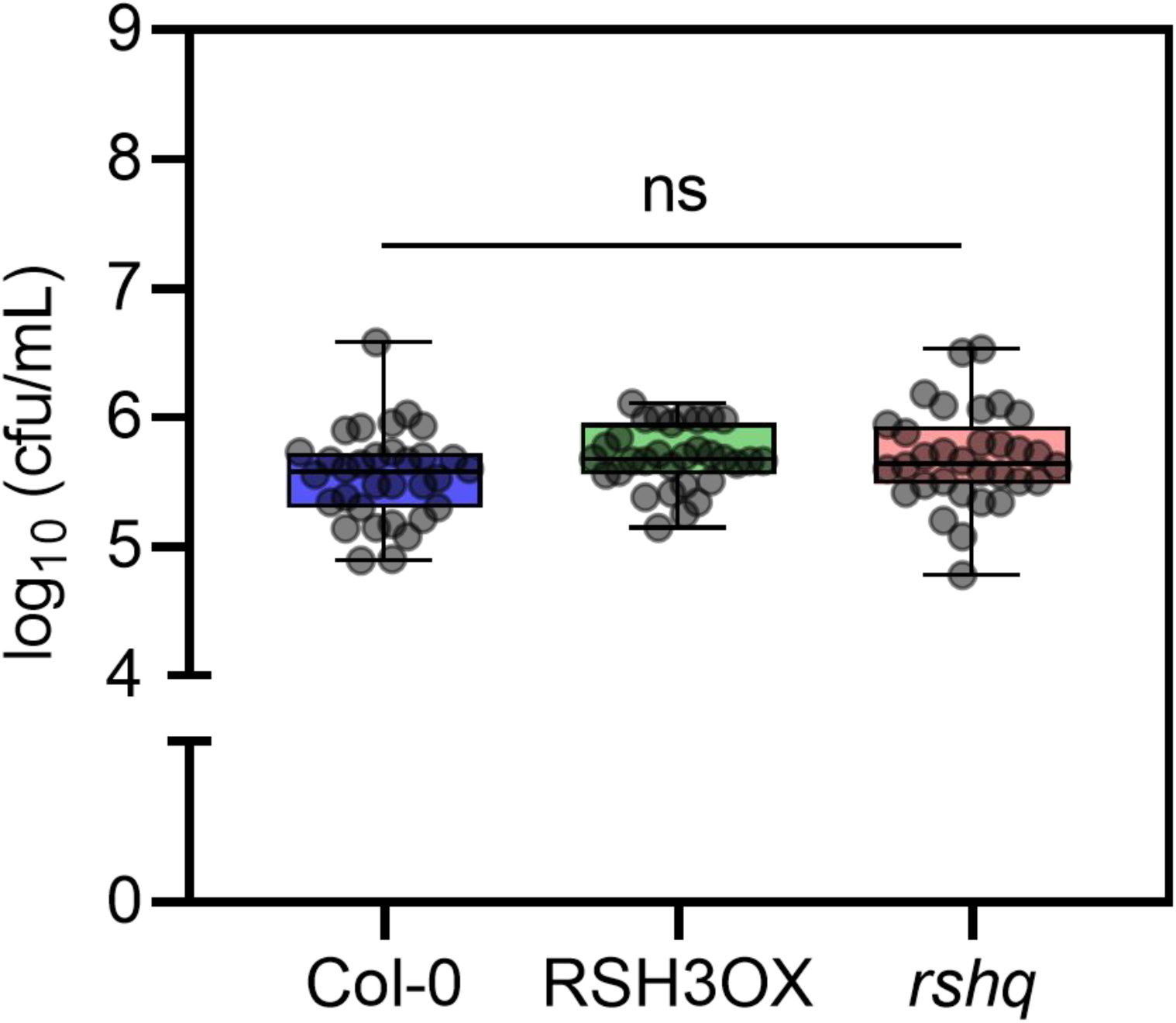
ppGpp levels do not affect the growth of a Δ*hrcC Pst* strain. Bacterial growth quantification in leaves of Col-0, RSH3OX, and *rshq* plants at 72h after inoculation with the type III secretion-deficient *Pst* mutant Δ*hrcC* (OD₆₀₀=0.1) Bacterial titers are expressed as log₁₀ (cfu/mL). ns: no significant difference among genotypes (one-way ANOVA with post-hoc test).

**Supplementary Figure S7.**
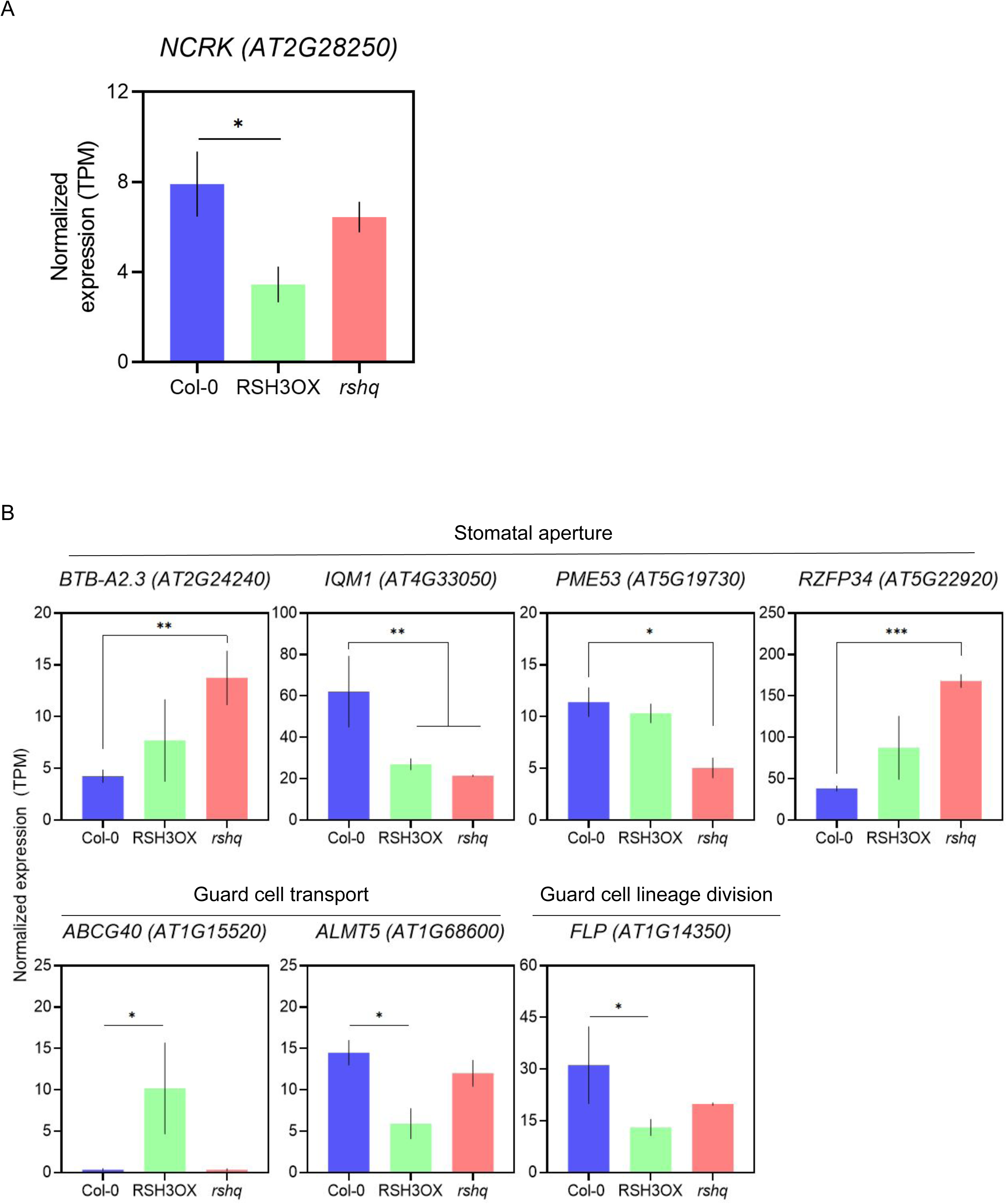
Expression of callose- and stomata-related (PTI) genes derived from RNA seq data. (A) Normalized expression (TPM) of *NCRK* from RNA seq experiment. Data are presented as mean ± SD. (B) Normalized expression (TPM) of *BROAD-COMPLEX, TRAMTRACK, AND BRIC-A-BRAC A2.3* (*BTB-A2.3*); *IQ-MOTIF CONTAINING PROTEIN1(IQM1*); *PECTIN METHYLESTERASE 53* (*PME53*); *ZINC RING-CONTAINING FINGER PROTEIN 34* (*RZFP34*); *ABC TRANSPORTER G FAMILY MEMBER 40* (*ABCG40*); *ALUMINUM-ACTIVATED MALATE TRANSPORTER 5* (*ALMT5*) and *FOUR LIPS(FLP*)/*MYB124* from RNA seq experiment. Data are presented as mean ± SD. Asterisks indicate significative differences among genotypes (adjusted p-value < 0.05).

**Supplementary Figure S8.**
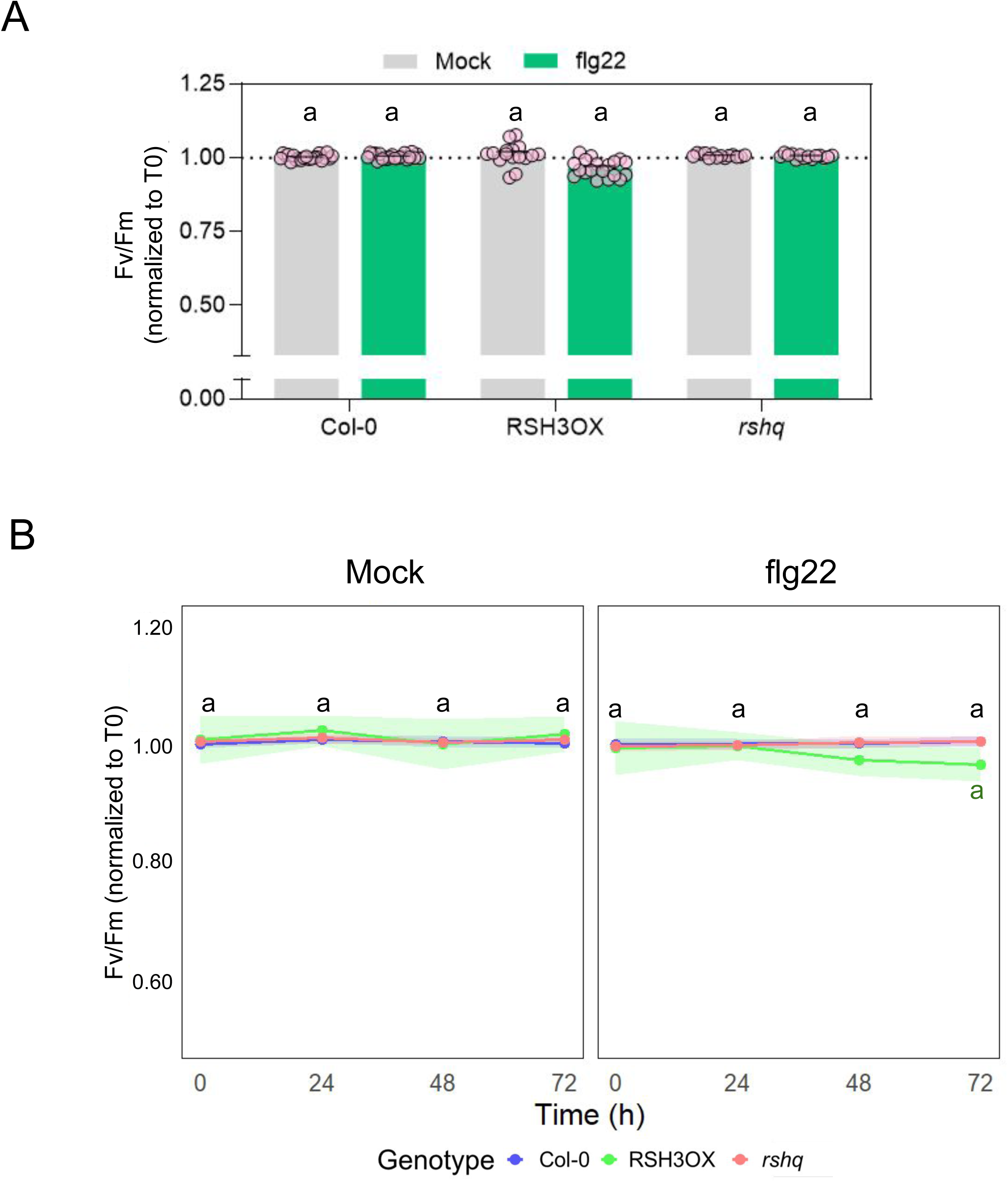
flg22 treatment does not affect photosynthetic efficiency. **(A)** Photosynthesis efficiency (Fv/Fm) of adult *A. thaliana* plants inoculated with H_2_O (mock) or flg22 1 μM. Data are shown in arbitrary units (a.u.) relative to 0h post-infection (T0). Data are presented as mean ± SEM. Different letters indicate statistical significant differences among genotypes and treatments (p<0.05, two-way ANOVA with post-hoc test). **(B)** Time course analysis of Fv/Fm during 72h for panel (A). Data are shown in arbitrary units (a.u) relative to 0h post-inoculation (T0). Lineplots are presented as mean ± SEM.

